# Synchronization and resilience in the Kuramoto white matter network model with adaptive state-dependent delays

**DOI:** 10.1101/2020.08.25.266841

**Authors:** Seong Hyun Park, Jeremie Lefebvre

## Abstract

White matter pathways form a complex network of myelinated axons that regulate signal transmission in the nervous system and play a key role in behaviour and cognition. Recent evidence reveals that white matter networks are adaptive and that myelin remodels itself in an activity-dependent way, during both developmental stages and later on through behaviour and learning. As a result, axonal conduction delays continuously adjust in order to regulate the timing of neural signals propagating between different brain areas. This delay plasticity mechanism has yet to be integrated in computational neural models, where conduction delays are oftentimes constant or simply ignored. As a first approach to adaptive white matter remodelling, we modified the canonical Kuramoto model by enabling all connections with adaptive, phase-dependent delays. We analyzed the equilibria and stability of this system, and applied our results to two oscillator and large dimensional networks. Our joint mathematical and numerical analysis demonstrates that plastic delays act as a stabilizing mechanism promoting the network’s ability to maintain synchronous activity. Our work also shows that global synchronization is more resilient to perturbations and injury towards network architecture. Our results provide key insights about the analysis and potential significance of activity-dependent myelination in large-scale brain synchrony.

## 1 Introduction

Synchronization, the mechanism by which oscillatory processes collectively organize to align their phase, has been the focus of intense research across the field of non-linear dynamics [1], especially in the biological sciences due to its involvement in myriads of physiological processes. In the brain, such synchronized oscillatory patterns, observed in the rhythmic discharge of neuronal electrical impulses, play a central role in neural communication, information processing, and the implementation of higher cognitive function [2]. However, the mechanisms by which these oscillations emerge and interact within and across brain microcircuits and systems remain poorly understood.

In the mammalian brain, local synchronized oscillatory activity are coordinated through a vast network of myelinated axonal fibers called white matter. The intricate white matter structure is formed under a population of glial cells called oligodendrocytes. The oligodendrocytes produce an insulating myelin sheath coiling around axonal membranes to greatly increase the conduction velocities of propagating signals between neurons. As white matter develops to adopt a genetically programmed structure, oligodendrocytes determine to which extent specific axons are myelinated. The resulting myelin structure of the network is responsible for the emergence and evolution of a rich repertoire of spatiotemporal activity patterns, notably oscillations with various spectral properties [3].

While white matter is highly relevant in shaping brain network dynamics, the mechanisms governing myelin development and its relationship with neural activity remain mostly unknown [4]. Nevertheless, the general hypothesis surrounding this topic has been shifting away from traditional viewpoints. Recent studies suggest that white matter structure undergoes continuous changes past the stages of developmental myelination and well into adulthood, rather than remaining static as initially presumed [5]. Indeed, substantial myelin formation continues to occur within the fully mature Central Nervous System in an activity-dependent manner [6, 7]. Furthermore, newly discovered evidence implies that white matter structure is responsive to experiences such as learning and social interactions [8]. These findings unveil potential ways white matter can restructure itself to facilitate neurological function over time. In particular, white matter characterized as a plastic, adaptive structure, can be critical in maintaining the essential oscillatory and synchronous processes found in many neural systems [9].

Despite the complexity of the neurophysiological processes involved, it has been hypothesized that myelin remodelling collectively reinforces synchronous dynamics and oscillatory coordination in large-scale brain networks [8]. From this perspective, the temporal structure of these plastic changes can provide a higher level of corrective adjustments, improving the network’s ability to converge towards phase synchrony. As a first approach to this intricate problem, we wish to use a simplified model to determine whether activity-dependent delays impact the phase coordination of coupled oscillators when compared to static delays. We have used the Kuramoto model, a phenomenological model which has a long history of applications in neuroscience, notably to study the collective organization of oscillatory neural systems [10]. The Kuramoto model, with or without time delay, has been extensively studied [1, 11, 12], and is increasingly used to model local oscillatory neural activity in brain-scale simulations informed by anatomical white matter connectivity [13, 14, 15]. Appealing for its relative simplicity, we have enabled the Kuramoto model with phase-dependent delays to examine the collective impact of adaptive conduction on coupled phase oscillators. Specifically, we performed the stability analysis of this modified system and examined its response to structural perturbations. While preliminary, these results can provide new insights into the large-scale impact of white matter remodelling and its potential role in neural synchrony.

This paper is structured as follows. In Section 2, we first introduce our network model composed as a system of coupled phase-oscillators with state-dependent delays. In Section 3, we derive the equations for an *N*-oscillator network’s synchronous state and its respective stability criteria. Section 4 applies the derived equations from Section 3 to a reduced two-oscillator network. Section 5 applies the derived equations from Section 3 to a large-dimensional oscillator network through taking an N-limit approximation by handling the asymptotic phases statistically. In Section 6, we conduct numerical experiments, and provide evidence supporting the analysis in Sections 4, 5 through simulated results. Section 7 explores the stabilization property in the context of spontaneous changes in network connectivity and compares the resilience of the synchronized state with and without plastic delays.

## 2 Model

We consider a prototypical model involving a network of *N* weakly-coupled nonlinear and delayed Kuramoto oscillators [9] whose phases *θ_i_*(*t*) evolve according to the following system of differential equations

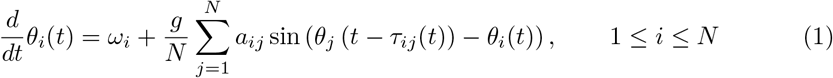

where 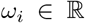 is the natural frequency of oscillator *i*, and *g* > 0 is a constant global coupling gain. The coefficients *a_ij_* represent synaptic weights: *a_ij_* = 1 if there is a white matter connection between oscillator *j* to *i*; otherwise, *a_ij_* = 0. The axonal conduction delays *τ_ij_* correspond to the conduction time taken by an action potential along a myelinated fiber. The propagation speed *c_ij_* = *c_ij_* (*t, ℓ*) fluctuates with respect to a propagating signal’s position *ℓ* along the length of an axon as saltatory conduction occurs at successive nodes of Ranvier. The conduction velocity *c_ij_* (*t, ℓ*) also changes in time. As such, each conduction delay may be written as

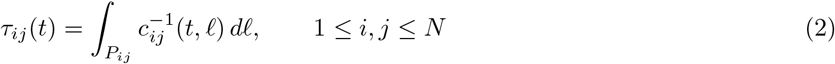

That is, a line integral along the axonal pathway *P_ij_* connecting oscillator *j* to *i*. It is uncertain how neural activity and/or oscillatory brain dynamics influences myelin formation. However, it is known that myelination correlates with information transmission within and across brain areas [5], which are known to involve long range synchronization [16]. To represent this in our model, we echo previous work on oscillatory neural communication [8, 9, 17] and make axonal conduction delays phase-dependent. Specifically, we assume that each conduction delay *τ_ij_*(*t*) evolves under the following sublinear plasticity equation

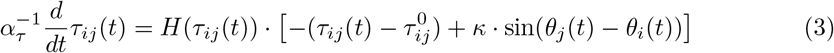

where 1 ≤ *i, j* ≤ *N*. Here, *α_τ_* is the homeostatic rate constant that sets the time scale of the evolving delays. The plasticity coefficient *κ* > 0 sets the gain of the conduction delay changes. The initial condition for delays *τ_ij_*(*t*) at *t* = 0 is provided by baseline lags 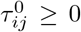. That is, 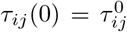 for all *i, j*. The Heaviside function *H*(*τ*) is defined as a smooth function such that *H*(*τ*) = 0 if *τ* < 0 and *H*(*τ*) = 1 if *τ* > *ε* for some small *ε* > 0. The Heaviside function *H*(*τ*) is present to preserve non-negativity of the delays *τ_ij_*(*t*) at all times *t* ≥ 0. For details on the construction of the smooth Heaviside function, we refer the reader to Appendix A. According to this rule, fluctuations in conduction delays are governed by an interplay between a local drift component that represents metabolic inertia towards an initial baseline lag 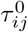 representing minimal myelination, and a forcing term that depends on the phase difference Δ_*ij*_:= *θ_j_* – *θ_i_* between oscillators *i* and *j*.

A schematic of the network structure as well as the activity-dependent plasticity rule are plotted in Fig. 1. There, given |*θ_j_* – *θ_i_*| < *π*, one can see that whenever *θ_j_* > *θ_i_*, the time delay increases due to an effective reduction in conduction velocity caused by myelin retraction. The opposite occurs if *θ_j_* < *θ_i_* and the time delay decreases: the connection speeds up due to myelin stabilization [8]. If the phase is the same, no changes in conduction velocity are required, and the delay remains stable at its initial lag 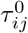.

**Figure 1.**
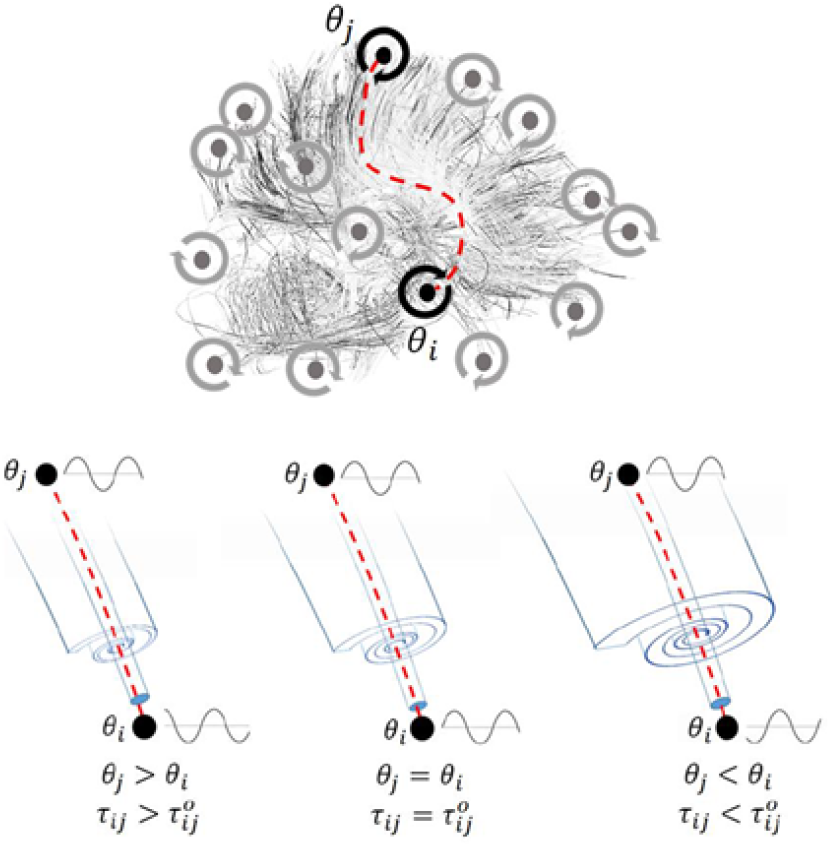
Schematic of a network of coupled oscillators with plastic conduction delays implementing temporal control of global synchronization. The network is built of recurrently connected phase oscillators, and connections are subjected to conduction delays. Those delays are regulated by a phase-dependent plasticity rule. Depending on whether the phase difference between two oscillators is positive, zero, or negative, the delays are either increased (myelin retraction), unaltered, or decreased (stabilization).

## 3 Synchronized state and stability with plastic delays

We are interested in characterizing the influence of the delay plasticity rule Eq. (3) towards the stability of global phase locked solutions and, more generally, in stabilizing synchrony between neural populations (i.e. oscillators). Enabled with adaptive delays, our model corresponds to an *N* + *N*^2^ dimensional functional differential equation with state-dependent delays. The analysis of such systems is technically challenging, especially in terms of stability where a modified approach must be used [18, 19, 20]. Mathematically, we analyze the network’s ability to asymptotically achieve the following synchronized state

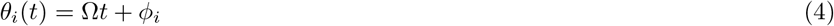

for all *i* ≤ *N* as *t* → ∞, where Ω is the global fixed frequency of the oscillators and *ϕ_i_* is the asymptotic phase-offset of oscillator i as time *t* → ∞. As we will see, adaptive delays following plasticity rule Eq. (3) require non-zero phase differences Δ_*ij*_(*t*):= *θ_j_*(*t*) – *θ_i_*(*t*) in order to maintain its equilibrium values, from which the network can no longer become in-phase. Nevertheless, the oscillators are able to become entrained to a common frequency Ω. Hence, we say that our network becomes synchronized when the nodes *θ_i_*(*t*) globally oscillate at some stable frequency Ω and become phase-locked under individual offsets *ϕ_i_*. As we are primarily concerned with the effects of delays changes under the plastic Eq. (3), we assume that all oscillators share the same natural frequency *ω_i_* = *ω*_0_ for all i in Eq. (1). To simplify the convergent behaviour of delays *τ_ij_*(*t*) following Eq. (3), we set *α_τ_* = 1.

Applying the ansatz Eq. (4) onto both the phases in Eq. (1) and the delays in Eq. (3), we obtain the following expressions for the global frequency 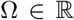 and individual offsets 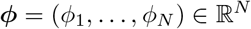 as

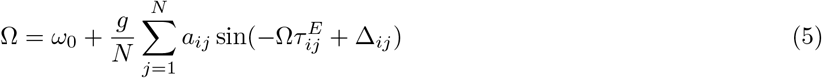

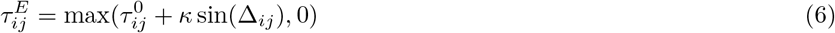

for all *i, j*, where Δ_*ij*_ = *ϕ_j_* – *ϕ_i_*. It must hold that if Ω satisfies Eq. (5), then Ω ∈ [*ω*_0_ – *g, ω*_0_+*g*]. To assess the network’s stability at the synchronized state (Ω, *ϕ*) satisfying Eqs. (5, 6), we introduce the perturbation terms *ϵ_i_*(*t*), *η_ij_*(*t*) around the equilibrium states Eq. (4) and Eq. (6) for *θ_i_*(*t*) and *τ_ij_*(*t*) respectively. That is, we write

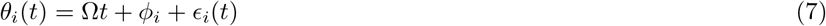

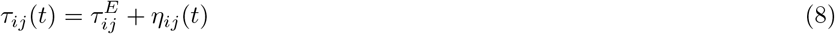

and examine the stability at the equilibrium point *ϵ_j_*(*t*) = *η_ij_*(*t*) = 0. The delay perturbations *η_ij_*(*t*) abide by the linearized form of Eq. (3) around all positive delays 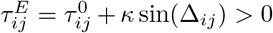. For the equilibrium delays 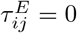, we proceed by assuming the corresponding perturbation terms will act in an asymptotically stable manner. Specifically, for all phase offset differences such that 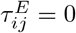, by the nature of the Heaviside cutoff function *H*(*τ*) in Eq. (3) there exists some 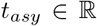 such that for all *t* > *t_asy_, τ_ij_*(*t*) = 0 and consequently *η_ij_*(*t*) = 0. That is, if 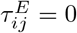, the perturbation term is asymptotically *η_ij_*(*t*) = 0. This condition will be satisfied given the synchronous state (Ω, *ϕ*) is indeed stable. Since the purpose of linearization is to determine stability, the above assumption is valid to make before proceeding. Taken together, the linearized equations for the terms *η_ij_*(*t*) as 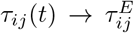 are given by

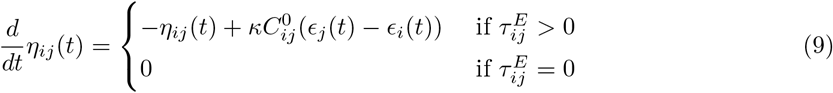

where 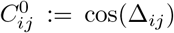. We can use the linearization approach in the context of equations with state dependent delays [18, 19, 20] as follows. Inserting Eq. (7) and Eq. (8) into the Kuramoto Eq. (1),

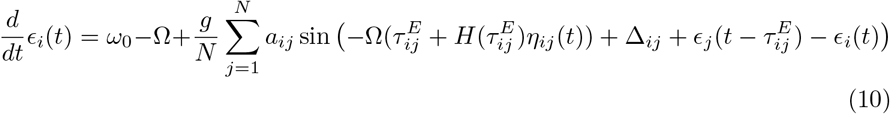

where we have made the approximation 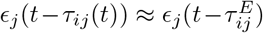 when *τ_ij_*(*t*) is near 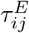. With (Ω, *ϕ*) satisfying Eq. (5), and by taking a first-order expansion around the term 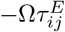, we obtain the linear system for the phase perturbation term given by

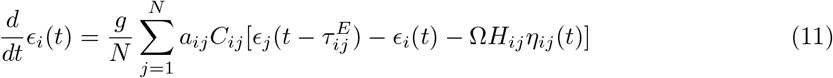

where we denote 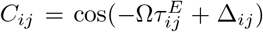 and 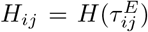. Hence, the network synchronizes at (Ω, *ϕ*) given that all perturbation terms *ϵ_i_*(*t*), *η_ij_*(*t*) following the linear system of fixed delay differential Eqs. (9, 11) converges to 0. From here, we can analyze the stability at the synchronous point (Ω, *ϕ*) by setting the ansatz 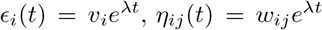 with respect to eigenvalue 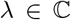. By Eq. (9), the coefficients *w_ij_* satisfy

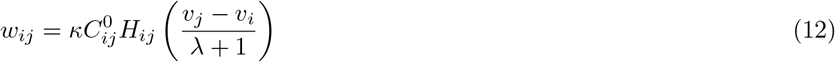

for all *i, j*, λ = –1. Applying Eq. (12) to coefficients *w_ij_*,

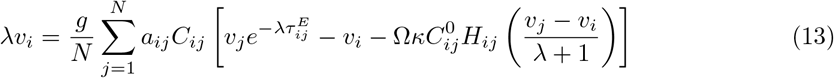

That is, λ is an eigenvalue if there exists an eigenvector 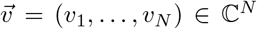 satisfying Eq. (13). In matrix form, Eq. (13) can be expressed as 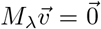, where *M*_λ_ = (*M_ij_*) is the N-dimensional square matrix with entries

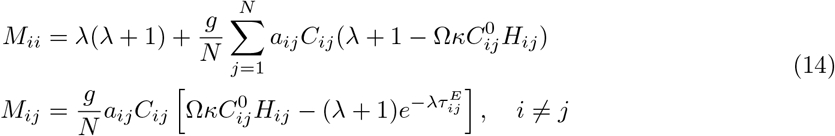

In summary, we acquire the following stability criterion: The system is stable around the synchronized state (Ω, *ϕ*) if for all eigenvalues 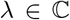 satisfying det *M*_λ_ = 0, Re(λ) < 0. Appendix A discusses the existence and uniqueness of solutions *θ_i_*(*t*), *τ_ij_*(*t*), as well as the justification of the above linearization and respective stability analysis.

Of note is that our approach is an extension of the non-plastic case. Indeed, without plasticity *κ* = 0, the delays remain fixed at the baseline lag 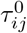, and as a result there is no need for the delays to establish an equilibrium with positive phase offset differences Δ_*ij*_. Hence, the oscillators become perfectly in-phase with *ϕ_i_* = 0 during synchronization. Consequently, Eq. (5) for the global frequency Ω reduces to the one-dimensional fixed-point expression

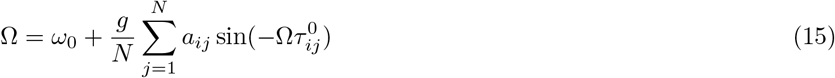

By employing similar steps as above, the phase perturbation terms *ϵ_i_*(*t*) follow the linear system

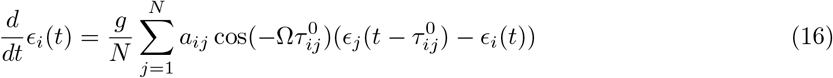

resulting in the corresponding eigenvalue matrix *M*_λ_ defined with entries

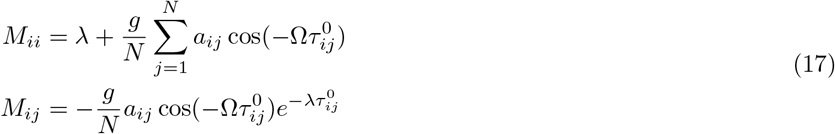

With non-plastic delays, stability analysis at *θ_i_*(*t*) = Ω*t* has been accomplished through other means. Lyapunov functionals [21] have shown that a sufficient criterion for synchronization around *θ_i_*(*t*) = Ω*t* is 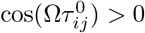 for all active connections *i, j* such that *a_ij_* = 1, and Ω satisfies Eq. (15). This criterion becomes necessary given a unique baseline lag 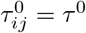 [22]. That is, it can be shown that the oscillators synchronize if and only if cos(Ω*τ*^0^) > 0.

## 4 Synchronization of two coupled oscillators with plastic delays

As a first illustrative approach to the general case when the number of coupled oscillators is large, let us first consider a reduced two-oscillator network. The following setup is similar to the system analyzed in [23]. We first let *N* = 2 with *a*_12_ = *a*_21_ = 1 and remove all self-integration terms by setting *a*_11_ = *a*_22_ = 0. We also set the baseline delays to be equal with 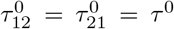. Then, the phases *θ*_1_(*t*), *θ*_2_(*t*) follow the Kuramoto system

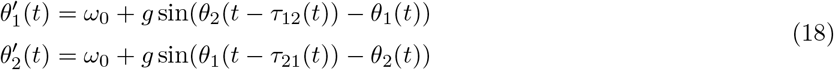

with respective plasticity equations

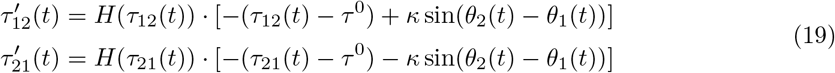

By symmetry we may let Δ_12_ = *ϕ*_2_ – *ϕ*_1_ > 0. We proceed by setting a sufficiently large plasticity gain with *κ* ≫ *τ*^0^. Then, it follows that the respective equilibrium delays are 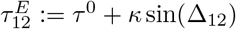 and 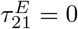, resulting in a single positive equilibrium delay. Hence, by Eq. (5) the frequency Ω and offset Δ_12_ satisfies

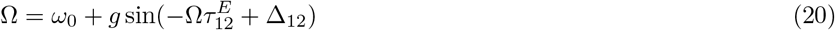

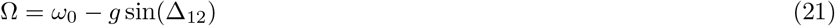

Solving for Δ_12_ above, we obtain

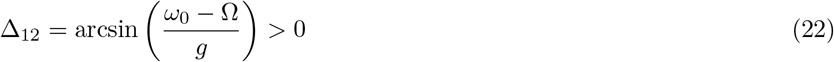

where positivity holds only when Ω < *ω*_0_. This leads to the oscillators synchronizing at a lower frequency Ω ∈ [*ω*_0_ – *g, ω*_0_). Substituting Δ_12_ in Eq. (20) with Eq. (22), if we define the root function

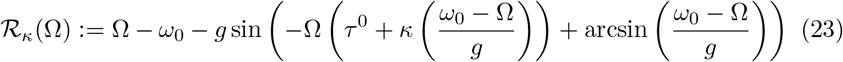

then *θ*_1_(*t*), *θ*_2_(*t*) synchronizes at a common frequency Ω implicitly satisfying *R_κ_*(Ω) = 0. Since it was assumed that 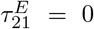, for self-consistency the synchronous frequency Ω must also satisfy τ^0^ – *κ* sin(Δ_12_) < 0, which can be written as Ω < *ω*_0_ – *gκ*^-1^ *τ*^0^ ≈ *ω*_0_ since we set *κ* ≫ *τ*^0^. Without plasticity *κ* = 0, the oscillators become in-phase with Δ_12_ = 0 and synchronizes at a frequency Ω that is a root of the function

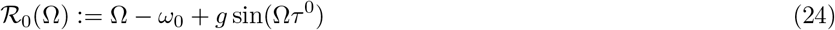

as stated by Eq. (15). Fig. 2(A) plots functions 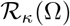 on the interval [*ω*_0_ – *g, ω*_0_) with respect to representative values *κ* = 0, 20, 30 of the plasticity gain. We observe that higher plasticity gains *κ* > 0 generally leads to a greater number of potential synchronization frequencies Ω for our system. In Fig. 2(A), one can see that the functions 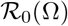 and 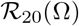 have single roots within the interval [*ω*_0_ – *g, ω*_0_), while 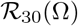 has five.

**Figure 2.**
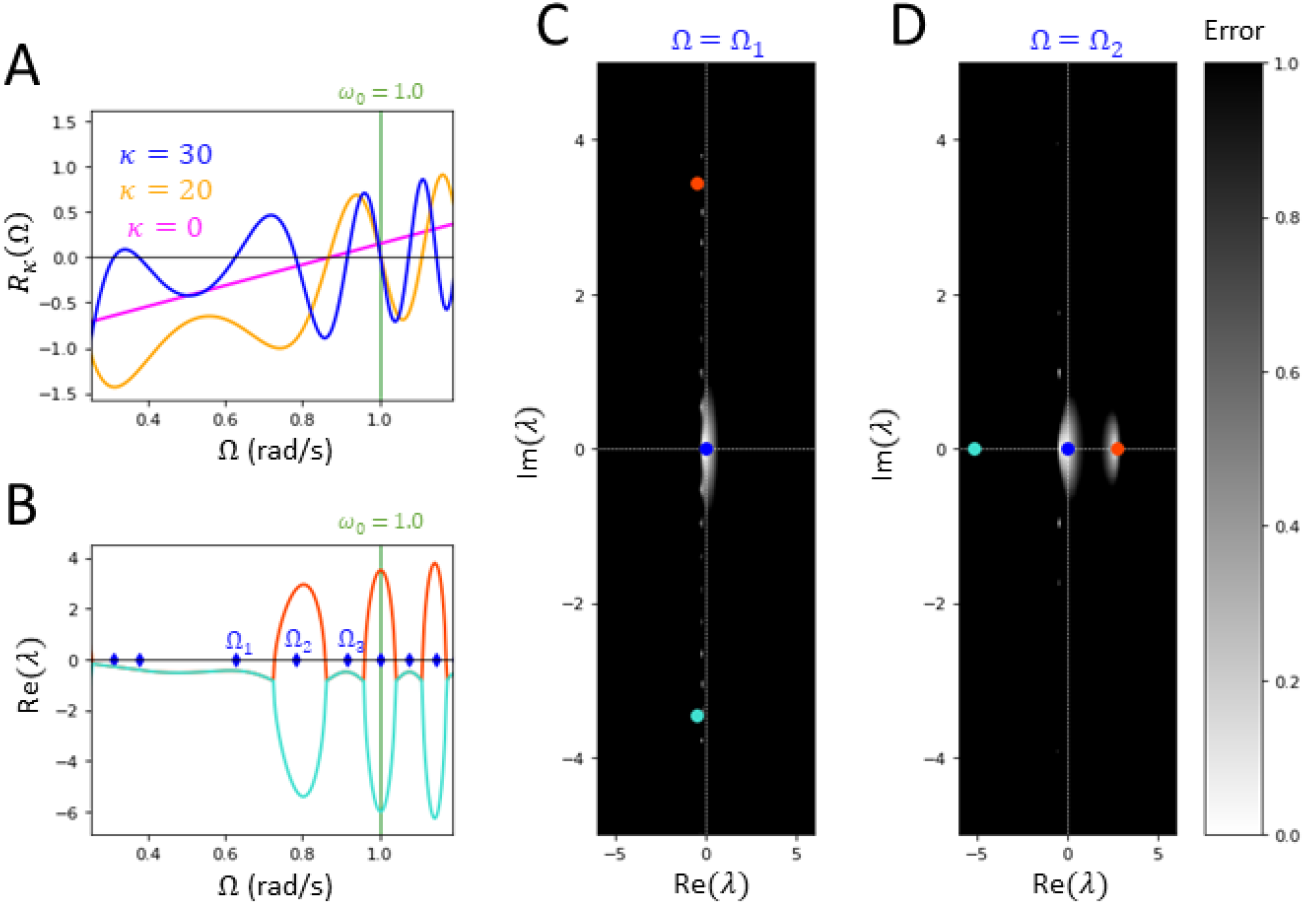
Theoretical stability plots for 2-oscillator system. A. Plot of error functions *R_κ_*(Ω) with varying fixed gain *κ* = 0 (magenta), *κ* = 20 (yellow), *κ* = 30 (blue). All roots Ω ∈ [*ω*_0_ – *g, ω*_0_) of *R_κ_* (Ω) are potential synchronization frequencies for the two oscillator system. The number of roots Ω for 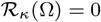 increase with larger *κ*. B. The plasticity gain is set to *κ* = 30. Plot of the real part of the non-zero branches λ_1_(Ω), λ_2_(Ω) (orange, cyan) of the polynomial root equation *P*_Ω_(λ) + *Q*_Ω_(λ) = 0 over Ω ∈ [*ω*_0_ – *g, ω*_0_). Ticks on the Ω-axis (blue) indicate the frequencies Ω_*i*_ solving 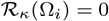 where the system can synchronize. The plotted branches imply that the oscillators will synchronize at Ω = Ω_1_, Ω_3_, and avoid the unstable frequency Ω = Ω_2_ with Re λ_1_(Ω_2_) > 0. C, D. Error heatmaps with Ω = Ω_1_, Ω_2_ respectively, that approximate the distribution of eigenvalues 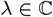 solving *P*_Ω_(λ) + *Q*_Ω_(λ)*e*^−λ*τ*^ = 0 near λ = 0, scaled and normalized for visibility. Spots near zero error (white) suggest potential eigenvalue locations. Markers plot the eigenvalues λ_0_ = 0, λ_1_ (Ω), λ_2_(Ω) (blue, orange, cyan) for *τ* = 0. The heatmap in D indicates an eigenvalue λ near λ_1_(Ω_2_) > 0, which implies instability at Ω = Ω_2_. All other eigenvalues λ appear to be distributed either at λ = 0 or on the left-side of the imaginary axis. Here, Ω_1_ = 0.626 and Ω_2_ = 0.783. For all plots, *α_τ_* = 0.5, *g* = 1.5/2, *ω*_0_ = 1.0, *κ* = 30, and *τ*^0^ = 0.1*s*.

The stability in the two-dimensional case can be easily determined. Indeed, as derived in Sect. (3), the stability of the oscillators at the synchronization state (Ω, Δ_12_) is determined by the distribution of eigenvalues 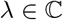 that satisfy

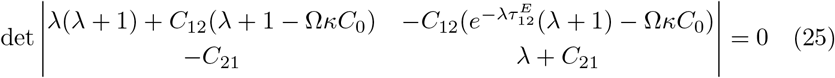

where 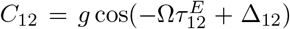, *C*_21_ = *g* cos(Δ_12_), *C*_0_ = cos(Δ_12_). This results in the root equation *P*_Ω_(λ) + *Q*_Ω_(λ)*e*^-λ*τ*^ = 0, where 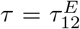 is the single positive delay and *P*_Ω_(λ), *Q*_Ω_(λ) are polynomials given by

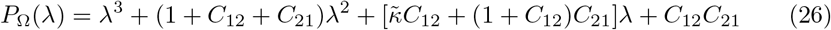

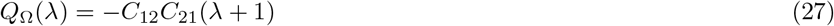

where 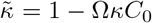. Note that *P*_Ω_(0) + *Q*_Ω_(0) = 0, which means we have neutral stability. If we make the approximation *τ* ≈ 0, the eigenvalues λ correspond to exact cubic roots of *P*_Ω_(λ) + *Q*_Ω_(λ). It is possible for the stability of the system under *τ* > 0 to align with the stability under *τ* = 0, particularly for small *τ*. If we ignore the neutral stability of our zero transcendental equation, Theorem 1 of [24] states that under certain conditions, the stability of the system at *τ* = 0 and at any *τ* > 0 does not change. For rigorous purposes however, we are still interested in finding the distribution of eigenvalues 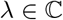 solving Eq. (25), particularly if it is accessible through numerical approximation.

Setting the plasticity gain *κ* = 30, Fig. 2(B) shows the real parts of the two non-zero branches λ_1_, λ_2_ of the roots *P*_Ω_(λ) + *Q*_Ω_(λ) = 0, plotted with respect to the global frequency Ω. If any of the branches are positive, it implies by the above discussion that the oscillators will not synchronize at the state (Ω, Δ_12_), where Δ_12_ = Δ_12_(Ω) is given by Eq. (22). We denote Ω_1_ < Ω_2_ < Ω_3_ to be the three largest synchronization frequencies corresponding to equilibria solutions Ω_*i*_ < *ω*_0_ of the root equation 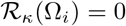 as presented in Fig. 2(A). The eigenvalue branches imply that frequencies Ω_1_, Ω_3_ are stable with Re(λ_1_), Re(λ_2_) < 0 at Ω = Ω_1_, Ω_3_, while Ω_2_ is an unstable frequency with Re(λ_1_) > 0 at Ω = Ω_2_. Fig. 2(C, D) shows heatmap approximations of complex roots λ of Eq. (24) at Ω = Ω_1_, Ω = Ω_2_ respectively. We can see that at Ω = Ω_1_, both non-zero cubic roots are located on the left-side of the imaginary axis, and the error heatmap shows that the eigenvalues are located on the left-hand side of the imaginary axis. At Ω = Ω_2_, all cubic roots are real with a single positive root λ_1_ > 0. Consistent with the stability of our system at *τ* = 0, the error heatmap implies there exists an eigenvalue 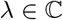 satisfying Eq. (25) near the positive root λ_1_. This highlights that under sufficiently large plasticity gain, adaptive delays introduce multiple stable states (Ω, Δ_12_) in a two-oscillator system, whose stability can be assessed under the zero delay approximation 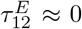. Our stability analysis reveals that the oscillators can synchronize at multiple possible frequencies, which suggests a greater degree of adaptability in our system.

## 5 Synchronization in large-scale oscillator networks with plastic delays

Using inspiration by the two dimensional case in Sect. 4, we consider here a large dimensional system using N-limit approximations. Again for simplicity, we set the baseline lags to be constant with 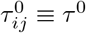 and the connection topology to be all-to-all with *a_ij_* ≡ 1. We approach the synchronization state (Ω, *ϕ*) of the network with adaptive delays, given by Eqs. (5, 6), in the following statistical sense. Suppose that the phase offsets *ϕ_i_* are i.i.d. under some distribution. We can set the offsets to be centered at 0 by defining the re-centered offsets 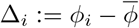, where 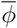 is the mean of *ϕ*. Then, Δ_*ij*_ = *ϕ_j_* – *ϕ_i_* = Δ_*j*_ – ̄_*i*_. We write that Δ_*i*_ are i.i.d. under some density function *ρ*(Δ). Setting a random Δ_*i*_ in the global frequency Eq. (5) for each *i* ≤ *N*, and then taking the limit *N* → ∞, we obtain the following *N*-limit approximation for frequency Ω and density *ρ*(Δ):

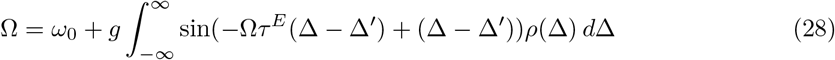

for all fixed 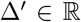, where *τ*^*E*^(Δ) = *H*(*τ*^0^ + *κ* sin Δ) · (*τ*^0^ + *κ* sin Δ). In order to apply Eq. (28) to obtain the global frequencies Ω, we parametrize the unknown density *ρ*(Δ) by assuming it is Gaussian under some small phase offset variance *δ*^2^. That is, we have

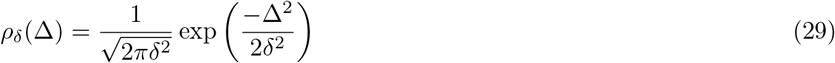

and we set *ρ*(Δ) = *ρδ*(Δ). If the variance *δ*^2^ is small, we can approximate the fixed offset in Eq. (28) as Δ′ ≈ 0, as well as make the approximation *τ^E^*(Δ) ≈ *H*(*τ*^0^ + *κ*Δ) · (*τ*^0^ + *κ*Δ) since Δ is small. Hence, our large-scale network synchronizes near a global frequency Ω with Gaussian distributed phase offsets with variance δ^2^ given that (Ω, *δ*^2^) is a root of the function

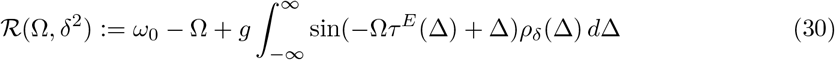

Without plasticity, we recall that we have in-phase synchronization at global frequency Ω which is a solution to 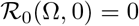. That is,

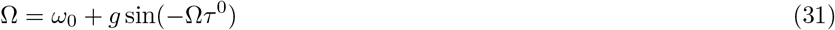

Fig. 3(A) plots the curve 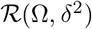 for various fixed-values of *δ* > 0 on Ω ∈ [*ω*_0_ – *g, ω*_0_ + *g*]. We find that there is a unique but different root Ω to 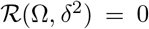 at each fixed variance *δ*^2^ > 0. Hence, we can obtain an implicit curve Ω = Ω(*δ*) by parametrizing the level curve 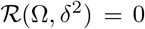 with respect to *δ* > 0. The curve is plotted in Fig. 3(B), and shows a continuous range of potential synchronization states (Ω(*δ*), *δ*^2^) along *δ* > 0 to a large *N*-dimensional system of oscillators. The graph of the level curve 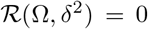 also shows that Ω(*δ*) < *ω*_0_ for all *δ* > 0, implying that the oscillators are drawn to synchronize at a lower frequency from their natural frequency.

**Figure 3.**
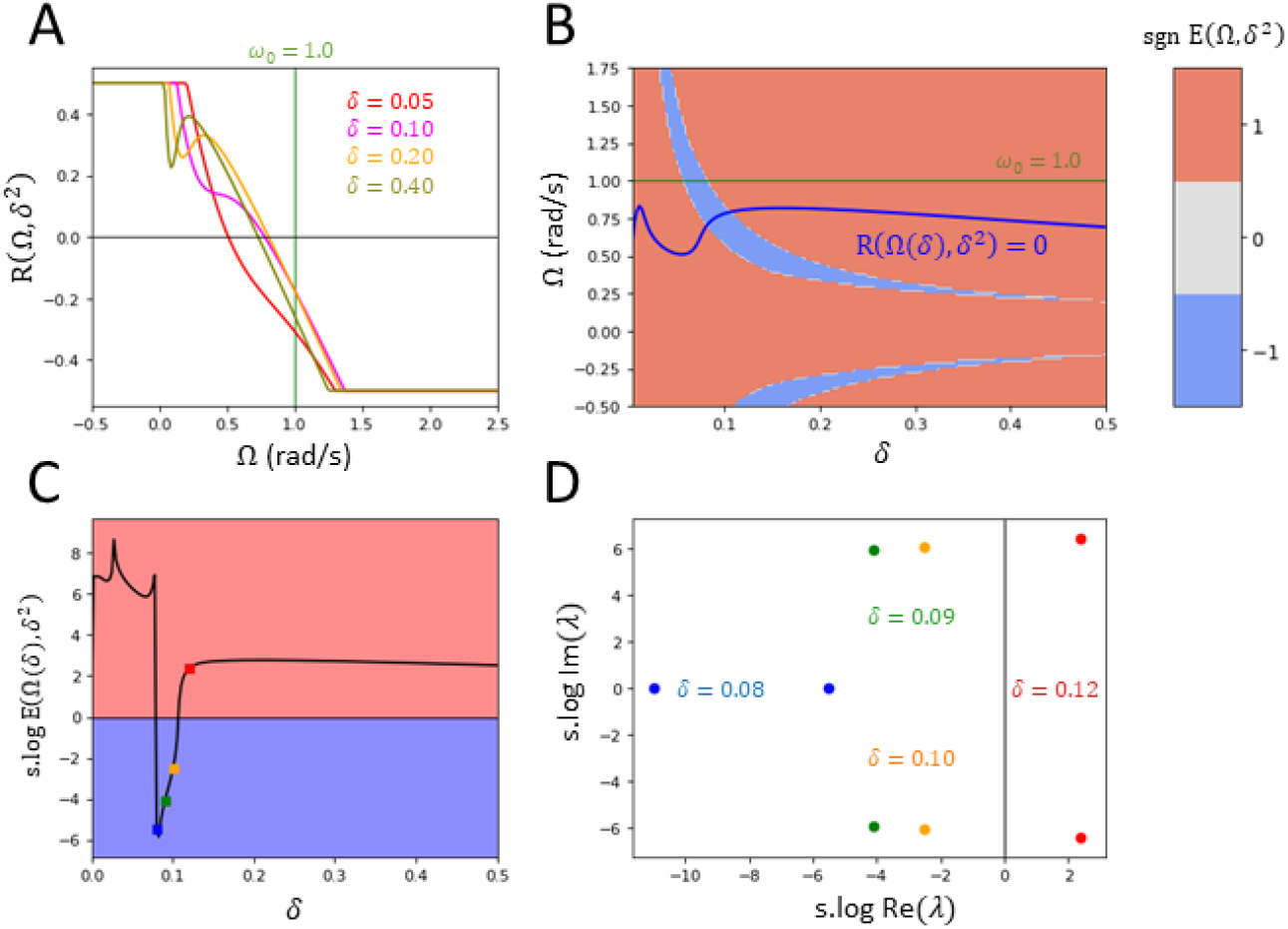
Theoretical stability plots for large N-dim oscillator system. A. Plots of error function 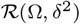 with varying fixed *δ* > 0 over Ω ∈ [*ω*_0_ – *g, ω*_0_ + *g*]. The function is truncated between interval [−0.5, 0.5] for visibility. There is a unique root 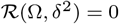 for each fixed *δ* > 0. B. Colour map of sgn *E*(Ω, *δ*^2^) over states (Ω, *δ*^2^) ∈ [*ω*_0_ – *g, ω*_0_ + *g*] × (0, 0.5^2^), along with the implicit solution curve (purple) Ω = Ω(*δ*) parametrizing level set 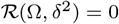. Stable regions correspond to sgn *E*(Ω, *δ*^2^) = −1 (blue) and unstable regions correspond to sgn *E*(Ω, *δ*^2^) = 1 (red). The network synchronizes near a state (Ω(*δ*), *δ*^2^) overlapping the stable region. C. Plot of stability term s. log *E*(Ω, *δ*^2^) along the solution curve Ω = Ω(*δ*) over *δ* ∈ (0,0.5). There is a small interval *δ* ∈ (0.08,0.1) for which (Ω(*δ*), *δ*^2^) is in the stable region (blue). Other states are in the unstable region (red). D. Complex plot of non-zero eigenvalues of *P*(λ | Ω, *δ*^2^) + *Q*(λ | Ω, *δ*^2^) on solution states (Ω(*δ*^2^), *δ*^2^) across varying *δ* > 0, scaled by s. log for visibility. The eigenvalues in plot D were computed at respective states (Ω, *δ*^2^) in plot C indicated by the same colour. Power terms for polynomial *Q*(λ | Ω, *δ*^2^) were computed up to degree *M* = 3. The parameters used for all plots are *α* = 1.0, *g* = 1.5, *ω*_0_ = 1.0, *κ* = 80, and *τ*^0^ = 0.1*s*.

For small offset differences, the equilibrium delays are approximately 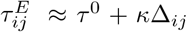 if Δ_*ij*_ > 0, and 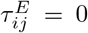 if Δ_*ij*_ < 0. Before proceeding, we set *κ* to be sufficiently larger than the baseline lag *τ*^0^ such that 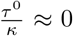. Then, most negative offset differences Δ_*ij*_ < 0 fall below the Heaviside cutoff *τ*^0^ + *κ*Δ_*ij*_ < 0. From this, the Heaviside term becomes approximately dependent to the sign of Δ_*ij*_ with *H_ij_* ≈ *H*(Δ_*ij*_) for all *i, j*. For large gain *κ*, two categories of equilibrium delays emerge that contribute to the global frequency Ω in Eq. (30): For half the connections with offset difference Δ_*ij*_ < 0, the corresponding delays *τ_ij_*(*t*) decay to 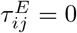 as we have *τ*^0^ < –*κ*Δ_*ij*_ < 0 with *τ*^0^ ≪ *κ*. Otherwise, with Δ_*ij*_ > 0 the delay *τ_ij_*(*t*) establishes a positive equilibrium at 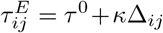. The positive delays 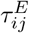 become widely distributed under large standard deviation *κδ*.

The coupled network can synchronize towards any stable point (Ω, *δ*^2^) along the curve *R*(Ω, *δ*^2^) = 0. To assess the stability at each state (Ω, *δ*^2^), consider our *N*-dimensional eigenvalue stability criterion 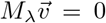 for eigenvector 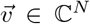 and matrix whose entries are given by Eq. (13). That is,

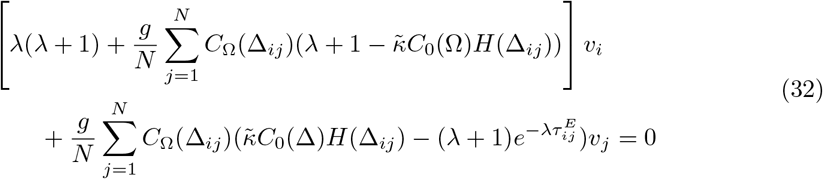

for all *i* ≤ *N*, where 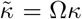, *C*_0_(Δ) = cos(Δ), *C*_Ω_(Δ) = cos(–Ω*t^E^*(Δ) + Δ), and *τ^E^*(Δ) = *H*(Δ)(*τ*^0^ + *κ*Δ). Once again, if our offset variance *δ*^2^ is small for Eq. (31) we can approximate the terms Δ_*ij*_ = Δ_*j*_ – Δ_*i*_ ≈ Δ_*j*_ by assuming Δ_*i*_ ≈ 0 at each *i*. We derive an *N*-limit version of the eigenvalue Eq. (32) as follows. Likewise to the frequency equation Ω, we can obtain a similar *N*-limit approximation to Eq. (32) as follows. We define a continuous eigenfunction 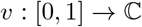 such that *v_i_* = *v*(*i/N*). Then, taking the limit *N* → ∞ to Eq. (32) with Δ_*ij*_ ≈ Δ_*j*_ ~ *N*(0, *δ*^2^), we obtain the continuous eigenvalue criterion for λ with respect to eigenfunction *v*(*x*) given by

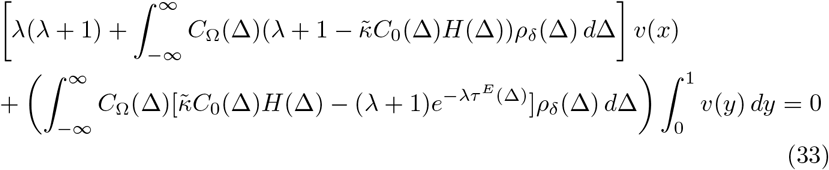

for every *x* ∈ [0,1]. Justifications regarding the *N*-limit step to derive Eq. (33) is provided in Appendix B. Here, the only continuous eigenfunction solution to the above equations is the constant function *v*(*x*) = 1^[1]^. Hence, Eq. (33) simplifies to the following N-limit eigenvalue equation for λ:

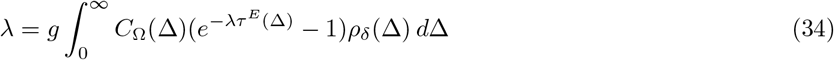

We claim that the the synchronization state (Ω, *δ*^2^) is (neutrally) stable given that for all 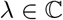 satisfying Eq. (34), Re(λ) ≤ 0. Note that for all λ = *u* + *iv* satisfying Eq. (34),

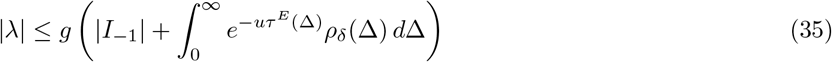

where 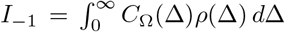. By Eq. (35), if *u* → ∞, |λ| ≤ *g*|I_-1_|, which is a contradiction. It follows that as |λ| → ∞, Re(λ) → –∞ for the distribution of eigenvalues λ satisfying Eq. (34). If *κ* = 0, then the frequency Ω solving Eq. (31) is (neutrally) stable if all eigenvalues 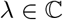 given by

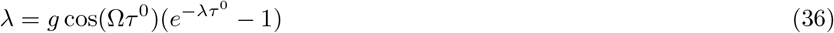

has non-positive real parts Re(λ) ≤ 0. As shown in [22], the non-plastic delay network synchronizes at Ω if and only if cos(Ω*τ*^0^) > 0.

At this point, the N-limit approximate eigenvalue Eq. (34) has done little to improve the N-dimensional criterion det *M*_λ_ = 0, due to exponential blow-up that persists within the integrand term. We proceed to reduce the eigenvalue Eq. (34) into an exponential polynomial root equation as follows. Rescaling λ → *R*λ by some radius *R* > *κ*,

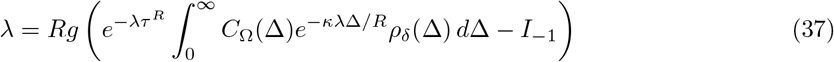

with rescaled small delay term *τ*^*R*^ = *τ*^0^/*R*. Expressing the power series of the exponential λ term up to degree *M* in Eq. (37) above, we obtain the approximate exponential polynomial root equation

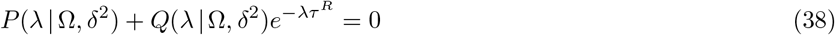

with polynomials

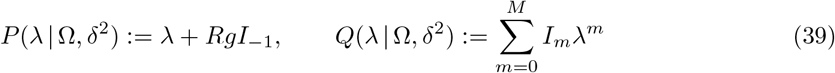

and power term coefficients

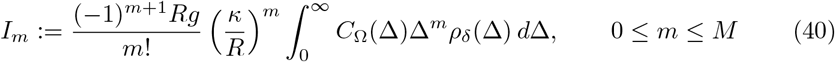

Choosing a large radius *R* > *κ*, the rescaled delay term *τ^R^* and coefficients *I_m_* for larger degrees *m* become arbitrarily small. From this, we claim that the stability at (Ω, *δ*^2^) is predominantly determined by the first few terms of the exponential power expansion. That is, the stability at synchronization state (Ω, *δ*^2^) is determined by the finitely many polynomial roots 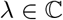 satisfying Eq. (38) with *τ^R^* ≈ 0 and some degree *M*. Denoting Λ(Ω, *δ*^2^) = {λ_1_,…, λ_*M*_} as the roots of *M*-degree polynomial *P*(λ | Ω, *δ*^2^)+*Q*(λ | Ω, *δ*^2^), the synchronization state (Ω, *δ*^2^) is (neutrally) stable given that

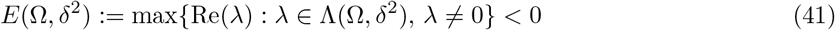

We note that these analytic results are reminiscent of what we obtained for twodimensional systems in Sect. 4 with the cubic exponential polynomial equation *P*_Ω_(λ) + *Q*_Ω_(λ)*e*^-λ*τ*^ = 0 as defined by Eqs. (26, 27).

As we experienced large scale fluctuations of stability term *E*(Ω, *δ*^2^), we processed its value with either the sign function sgn (*x*) or the sign logorithm s. log(*x*) defined as

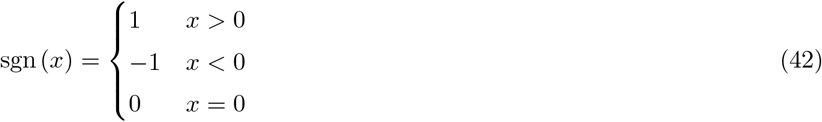

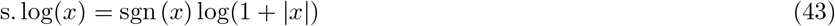

In Fig. 3(B), we plot sgn *E*(Ω, *δ*^2^) for all Ω ∈ [*ω*_0_ – *g, ω*_0_ + *g*] and small *δ* > 0. Stable regions are indicated where sgn *E*(Ω, *δ*^2^) = –1. In computing *E*(Ω, *δ*^2^), eigenvalues satisfying |Re(λ)| > 10^-8^ were considered to be non-zero. We notice that a section of the level curve 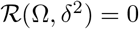 overlaps part of the stable region for (Ω, *δ*^2^). Indeed, Fig. 3(C) plots the stability term s.log *E*(Ω, *δ*^2^) along all points (Ω, *δ*^2^) on the implicit solution curve Ω = Ω(*δ*). As the plot shows, there is a small interval (*δ_l_, δ_r_*) such that (Ω(*δ*), *δ*) is stable for all *δ* ∈ (*δ_l_, δ_r_*) such that *E*(Ω, *δ*^2^) < 0. Fig 3(D) shows the transition of the eigenvalues 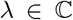 of *P*(λ | Ω, *δ*^2^) + *Q*(λ | Ω, *δ*^2^) = 0, corresponding to state (Ω(*δ*), *δ*^2^) as *δ* leaves the interval (*δ_l_, δ_r_*). We see that the eigenvalues shift towards the right-side of the imaginary axis, which shows the transition from stability to instability along the solution curve (Ω(*δ*), *δ*^2^).

The results above were obtained by setting the polynomial degree to *M* = 3 for *Q*(λ | Ω, *δ*^2^), while the approximation failed for powers *M* > 3. In addition, it remains unresolved whether the distribution of eigenvalues λ_*N*_ satisfying the *N*-dimensional Eq. (32) with some eigenvector 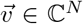 generally converge to the *N*-limit eigenvalues λ satisfying Eq. (33) with corresponding continuous eigenfunction 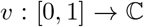. That is, whether for each *ϵ* > 0 there is some large *N* such that for all 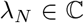 satisfying Eq. (32) each at *N* dimensions, there exists some eigenvalue 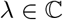 satisfying Eq. (33) such that |λ_*N*_ – λ| < *ϵ*. Integral equation theory has proven some relevant theorems. For instance, it can be proven [25] that there exists a nonzero eigenvalue 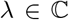 to Eq. (33) such that λ_*N*(*k*)_ → λ for some subsequence of eigenvalues 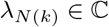 satisfying Eq. (32) at dimension *N* = *N*(*k*). Nevertheless, our *N*-limit analysis highlights an approximate region of points (Ω, *δ*^2^) where a large-dimensional system of oscillators will synchronize. We demonstrate in the following section that this region aligns with synchronous behaviour in numerical simulations.

## 6 Comparison to numerical simulations

Here, we validate the theoretical analysis committed in the previous sections through comparisons with numerical simulations. We obtain the global frequency Ω and asymptotic phase offsets φ¿ numerically, and systematically compare them with their analytical counterparts. Given numerical solution *θ_i_*(*t*), *i* ≤ *N* to the Kuramoto Eq. (1) with corresponding derivative solution 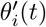, *i* ≤ *N*, we can obtain the asymptotic frequencies for each oscillator by summing over the time interval [*t*, *t* + *h*] and taking the limit *t* → ∞. That is,

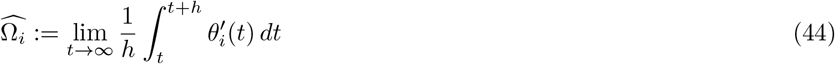

and estimate the global frequency Ω as the sample mean of individual frequencies

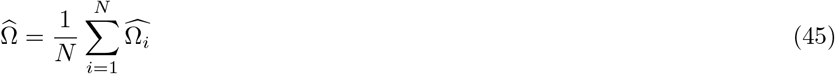

Likewise, we can numerically estimate the asymptotic phase offsets *ϕ_i_* for each oscillator by taking the difference 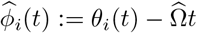 and defining the limit

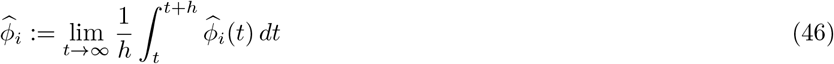

After modding 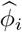 so that 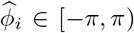, we can estimate the offset variance *δ*^2^ by taking the sample variance

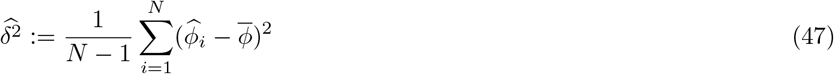

and the average (asymptotic) phase 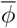 by taking the sample mean,

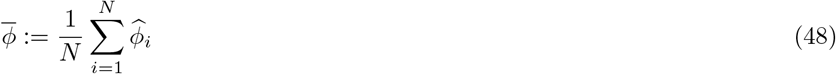

If our solutions *θ_i_*(*t*) synchronize towards some synchronous frequency Ω, offsets *ϕ_i_*, then 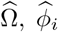 are estimators for Ω, *ϕ_i_* respectively. Numerically, we evaluate the estimators by taking the average over interval [*t, t* + *h*] starting at some large time *t* > 0.

We set all baseline delays to be a unique value *τ*^0^ > 0, so that 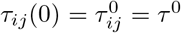. Since the plasticity rule Eq. (3) is an Ordinary Differential Equation, it suffices for all delays to have an initial value at *t* = 0. Before *t* = 0, we set the phases *θ_i_*(*t*) to be positioned in accordance to some initial frequency Ω_0_ and initial phase offsets 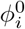. That is, we define the linear initial function 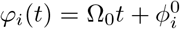 and set *θ_i_*(*t*) = *φ_i_*(*t*) on *t* ≤ 0^[2]^. In order to have reasonably behaving solutions *θ*(*t*), we modify the function *φ_i_*(*t*) so that it satisfies the necessary condition

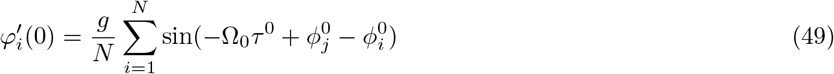

for all *i*. This adjustment is needed in order to avoid numerical discontinuities for 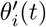. For details, refer to Appendix C.

Fig. 4 plots the results of a series of numerical simulations in the reduced two oscillator network as set up in Sect. 4. All trials were run from 0-200s, with estimated values 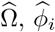 obtained by averaging the arrays over the last 20 seconds. The same parameter values used in Fig. 2 were applied here when running the numerical trials. We demonstrate the existence of two stable synchronization states with different frequencies. Fig. 4(A, B, C) shows the asymptotic behaviour of two trials (purple, orange) that started with different initial functions, plotting over the first 50 seconds. Fig. 4(A) plots the derivative arrays 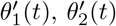 of each trial. We observe in Fig. 4(A) that each pair of oscillator frequencies entrain to a common frequency over time. The two trials converge to different common frequencies, estimated to be 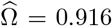 (purple lines) and 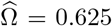 (orange lines) respectively. Fig. 4(B) plots the offset arrays 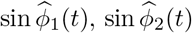 of each trial. For each trial, the two oscillators become phase locked as 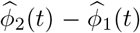 converges asymptotically to estimated constant differences 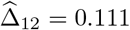 (purple lines) and 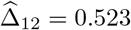 (orange lines), where 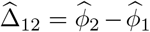. Fig. 4(C) plots the connection delays *τ*_12_(*t*), *τ*_21_(*t*) over time. By choosing our plasticity gain *κ* = 30 0.1 = *τ*^0^, it follows that for both trials, *τ*_12_(*t*) converges to some positive equilibrium delay *τ^E^* and *τ*_21_(*t*) decays to 0.

**Figure 4.**
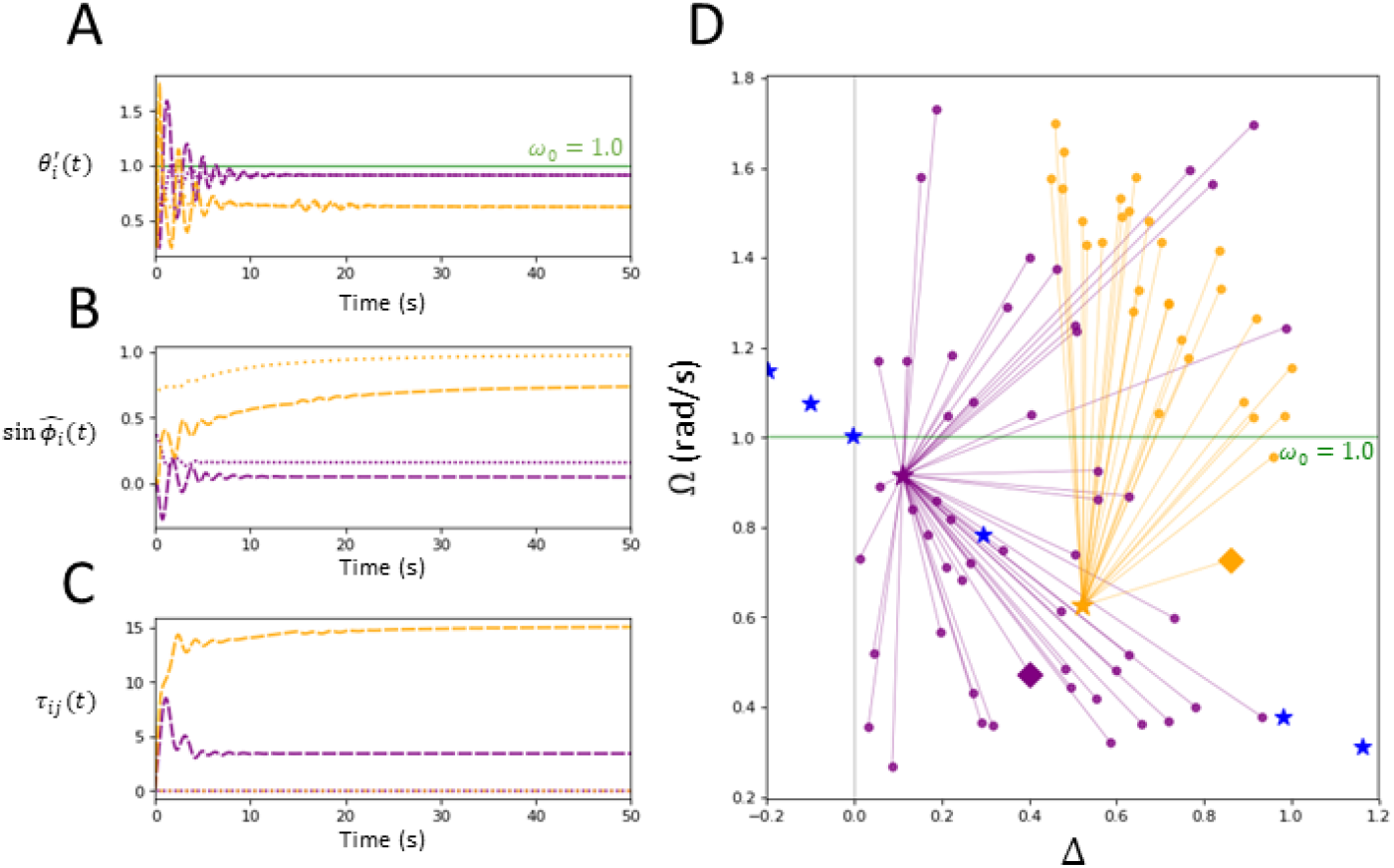
Numerical plots for 2-oscillator system. For plots A,B,C, two trials with different initial conditions are graphed. Trial 1, 2 (purple, orange) starts with initial frequency and phase difference (Ω_0_, Δ_0_) = (0.473,0.402), (0.727, 0.860) respectively. A. Plots of derivatives 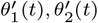 over time. For each trial, 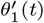 (dashed) and 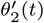 (dotted) converge to a common value 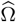 asymptotically. Each trial of oscillators entrain to a different frequency, implying the existence of multiple synchronization frequencies. B. Plots of sine phases 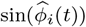, where 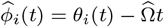. For both trials, the oscillators asymptotically phase-lock with 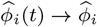, *i* =1 (dotted), *i* =2 (dashed). The phase-lock difference 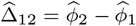 is also different for the two trials. C. Plots of adaptive delays *τ*_12_(*t*) (dashed), *τ*_21_(*t*) (dotted) over time. For each trial, delay *τ*_12_(*t*) converges to some positive equilibrium *τ^E^*, and delay *τ*_21_(*t*) decays to 0. D. Plots showing where two oscillators with randomized initial conditions (Ω_0_, Δ_0_) (orange) synchronize towards in terms of asymptotic frequency and phase offset 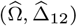 (magenta) across 80 trials. Each of the two trials in A, B, C have their initial condition plotted as a diamond marker of matching colour. Theoretical synchronization states (Ω, Δ_12_) given by Eqs. (20, 22) are also plotted (blue). Trials converge to two states 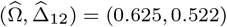, (0.916, 0.111), which align with the two theoretically stable states shown in Fig. 2. The parameters used for all plots are *α_τ_* = 0.5, *ε* = 0.01, *g* = 1.5/2, *κ* = 30, *ω*_0_ = 1.0, *τ*^0^ = 0.1*s*.

The above results imply that there exists at least two stable synchronization states, and that the frequency the system converges towards is dependent on the initial functions *φ*_1_(*t*), *φ*_2_(*t*). To provide further confirmation towards this proposition, the trials shown in Fig. 4 were repeated 80 times randomized across sampled initial frequency Ω_0_ ∈ [*ω*_0_ – *g, ω*_0_ + *g*] and initial phase difference Δ_0_ ∈ (0,1). The point (Ω_0_, Δ_0_) (circle markers) defines the initial functions *φ*_1_(*t*) = Ω_0_*t, φ*_2_(*t*) = Ω_0_*t* + Δ_0_. The theoretically stable states corresponding to frequencies Ω = Ω_1_ and Ω = Ω_3_ are plotted as orange and purple stars respectively, which are both close to the asymptotic frequency of the matching coloured trial discussed above. The rest of the theoretical synchronous states given by the roots of Eq. (23) are plotted as blue stars. Each trial’s solution arrays synchronized near one of the two stable states, as shown by the connecting coloured lines. The matching colours indicate which of the two stable frequencies Ω_*i*_ the trial’s solution arrays synchronized towards, such that the 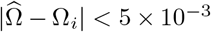. Each of the two trials graphed in Fig. 4(A, B, C) are also plotted in Fig. 4(D) with a diamond marker of matching colour. Convergence of the points is suggestive of a separatrix curve between the basins of attraction of both stable fixed points. It was also observed that the system synchronized towards the frequency Ω = Ω_3_ faster than Ω = Ω_1_, which suggests that the state Ω = Ω_3_ has a greater force of attraction. Hence, the experimental results align with the analysis outlined in Sect. 4. We draw the conclusion that in our reduced two-oscillator system, plastic delays are able to generate multiple synchronization states in comparison to non-plastic delays.

Fig. 5 provides the numerical results of a similar experiment performed as in Fig. 4 with a large dimensional network N = 50 and all-to-all network. All trials were run from 0-100s, with estimated values 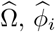 obtained by averaging the arrays over the last 10 seconds. The same parameter values are used as in Fig. 3. The initial function *φ*(*t*) for each trial was set up by choosing some frequency 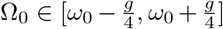 and deviation *δ*_0_ ∈ (0, 0.5). The initial phases 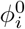 were i.i.d. sampled uniformly from the interval 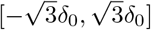. The *i*th oscillator was equipped with the initial linear function 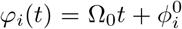. Fig. 5(A, B, C, D) graphs a single numerical trial with Ω_0_ = 0.913, *δ*_0_ = 0.295. Fig. 5(A) plots the derivative arrays 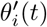, which we note converges to some constant frequency estimated as 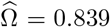. Fig. 5(B) plots the offset arrays sin 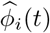, which shows that each 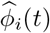 converges to some constant phase offset estimated by 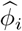. Hence, the oscillators become asymptotically phase-locked under distributed offsets with estimated variance 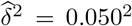. Fig. 5(C) plots a sample of 50 adaptive delays *τ_ij_*(*t*), which become part of the positive equilibrium distribution 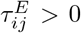 or decay to 0. Fig. 5(D) plots the density of centered phases 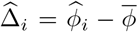. As we assumed that the centralized phase offsets follow a Gaussian distribution, we perform a normality test on the numerical asymptotic offsets 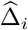. The Shapiro-Wilk Test for non-normality returned a p-value of 0.005 for 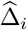, which suggests another distribution would be more accurate. Visually, a Gaussian curve 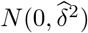 (black line) is fit over the density in Fig. 5(D). Nevertheless, the relevance of the Gaussian approximation, which greatly simplifies the analysis, becomes apparent as the numerical and analytical results nearly coincide. Other approximations could be used to facilitate the analysis further, and are left for future work.

**Figure 5.**
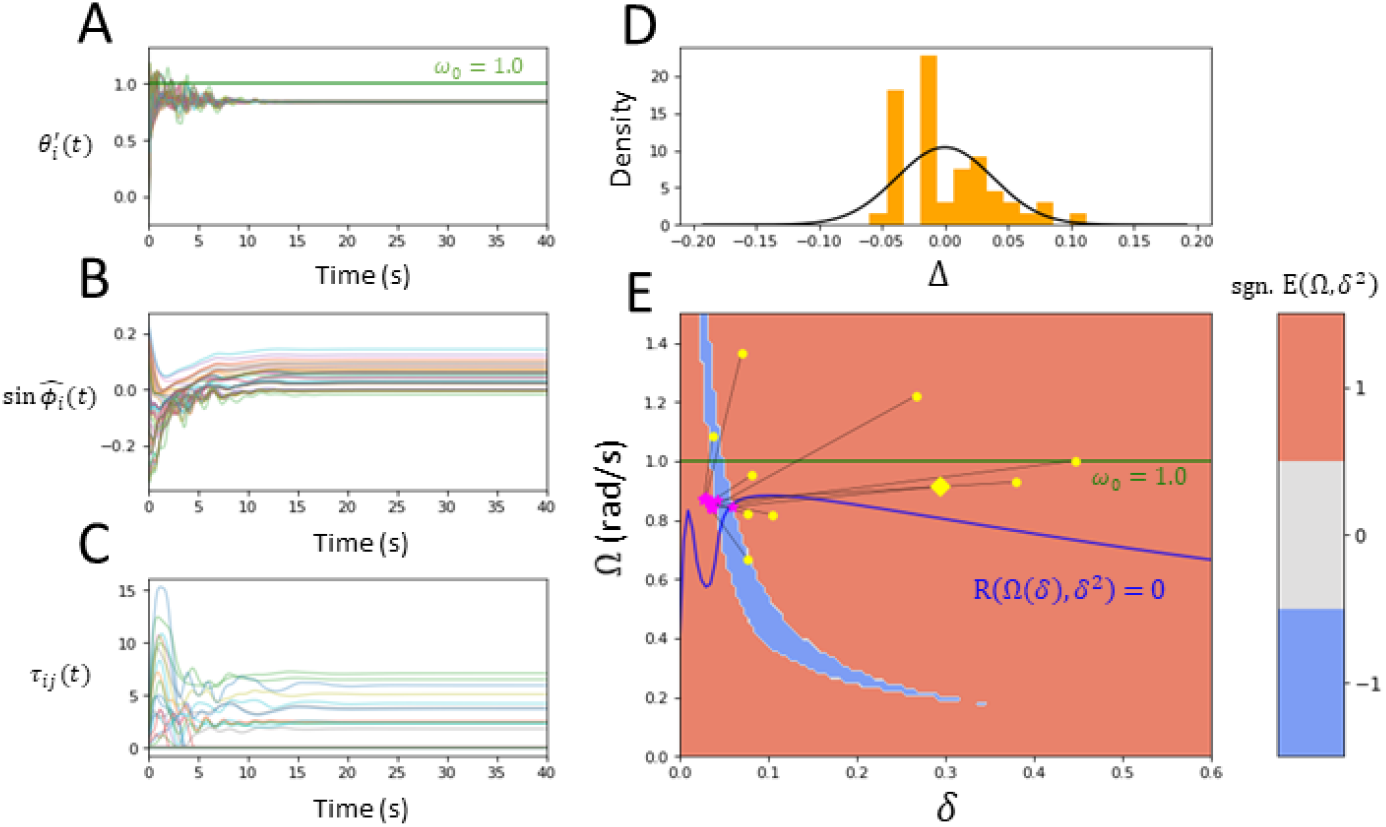
Numerical plots of N-oscillator system. A. Plots of derivatives 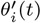 over time. We have that all 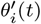 converge to a common frequency, estimated to be 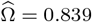. B. Plots of sine phases 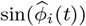, where 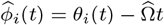. All oscillators appear to asymptotically phase lock to one another. C. Plots of a sample of 50 adaptive delays *τ_ij_*(*t*) over time. Some delays *τ_ij_*(*t*) converge to some positive equilibrium 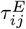, while others decay to 0. D. Density of centralized phase offsets 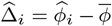, which was assumed to be Gaussian. The Gaussian curve 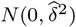 (black line) is fit over the density. E. Plot showing oscillators with randomized initial frequency and phase deviation (Ω_0_, *δ*_0_) (yellow) synchronizing towards respective estimated asymptotic frequency and phase deviation 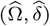 (magenta star) across 10 trials. The yellow diamond refers to the trial plotted in A,B,C. The numerical values are plotted directly over Fig. 3(B). As shown, the network synchronizes near the theoretical stable region. The parameters used were *N* = 50, *ε* = 0.01, *α_τ_* = 0.1, *g* = 1.5, *κ* = 80, *ω*_0_ = 1.0, *τ*^0^ = 0.1*s*.

The numerical simulation, as presented above, was repeated 10 times with randomized initial conditions 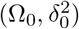. For each trial, Ω_0_, *δ*_0_ was sampled uniformly from intervals 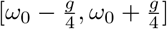 and (0,1) respectively. Fig. 5(E) plots the following convergence results. Each trial with initial condition 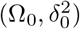 (yellow markers) synchronized near the respective estimated point 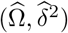 (magenta markers). We can see that every trial synchronized at approximately the same state (Ω, *δ*^2^). To determine whether the numerical results align with the analysis discussed in Sect. 5, the numerical values were plotted on top of Fig. 3(B). We observe that for each trial, the network synchronizes near the portion of the level curve 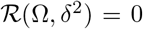 within the stable region where *E*(Ω, *δ*^2^) < 0. Hence, the numerical experiment for *N* = 50 generated results that validates the theoretical *N*-limit stability analysis in Sect. 5.

## 7 Neuroscience application: resilience to injury with sparse and uniform connectivity

One of the most salient examples of white matter plasticity comes from neuroimaging in presence of lesion. In these cases, white matter remodelling takes place in order to restore and maintain function, a process that notably impacts neural synchronization [26]. Let us now investigate whether the plasticity mechanism can be used to stabilize phase locked states in the presence of network damage. That is, we would like to know whether changes in time delays can be used to compensate for a reduction in effective connectivity and make the global synchronous state more resilient. To investigate this problem, we model injury as a loss in connections *a_ij_*. Defining *γ* ∈ [0,1] as the insult index, we introduce here the sparse synaptic connectivity weights given by

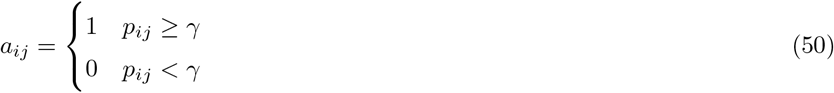

where *p_ij_* is a uniformly distributed i.i.d. sampling on [0,1]. *γ* represents the connectivity damage through an increase in the sparseness of network connections. Note that if *γ* = 0, then we obtain the all-to-all connection topology *a_ij_* ≡ 1 corresponding to no injury in the system. If *γ* = 1, we have the trivial system *a_ij_* = 0 which means no signals between the oscillators occur. For any *γ* we have the probability to retain the connection 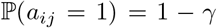. Without plasticity, the mean phase dynamics for *N* → ∞ is given by

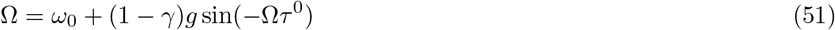

by Eq. (31). By observation, one can see that we are in the presence of the same network dynamics, but with an effective coupling coefficient given by *g*_eff_:= (1–*γ*)*g*. Thus, damage simply reduces the net coupling. To demonstrate this point, we start with a strong coupling parameter and decrease it until stability is either lost or preserved by plasticity. According to Eq. (51), Ω → *ω*_0_ as *γ* → 1. Hence, if the baseline lag *τ*^0^ is chosen such that cos(Ω*τ*^0^) ≥ 0 and cos(*ω*_0_*τ*^0^) < 0, the network without plastic delays is susceptible to injury destabilizing the synchronous state.

We ran numerical simulations by applying similar parameter values as Sect. 6 while introducing injury *γ* = 0.8 to the connections at time *t* = *t_inj_*, set at *t_inj_* = 80*s*. The initial condition of the network was fixed at (Ω_0_, *δ*_0_) = (*ω*_0_, 0.25) for all trials. Fig. 6(A) shows the destruction of existing connections following injury, comparing connection grids for *a_ij_* before and after *t_inj_*. Fig. 6(B) shows the distribution of the delays *τ_ij_*(*t*) with existing connections *a_ij_* = 1 at timestamps *t* = 0*s*, *t* = 79*s* (pre-injury), *t* = 160*s* (post-injury). With fewer connections available, the ability for the surviving delays to adjust themselves are crucial in re-stabilizing the system’s synchrony. Fig. 6(C) (no gain) and Fig. 6(D) (with gain) show that both networks entrain to a global frequency successfully before and after inflicted injury towards the network. The entrainment frequencies pre-injury 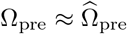 and post-injury 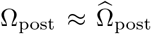 were estimated by taking the average of 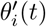 at times 148 – 160s and 304 – 320s respectively for each *i* ≤ *N*. Fig. 6(E) (no gain) and Fig. 6(F) (with gain) plots the sine phase offsets sin 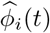 over time, given by

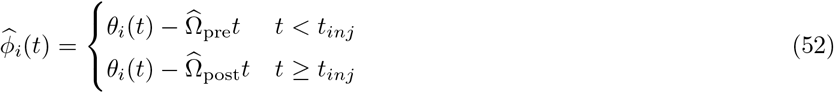

**Figure 6.**
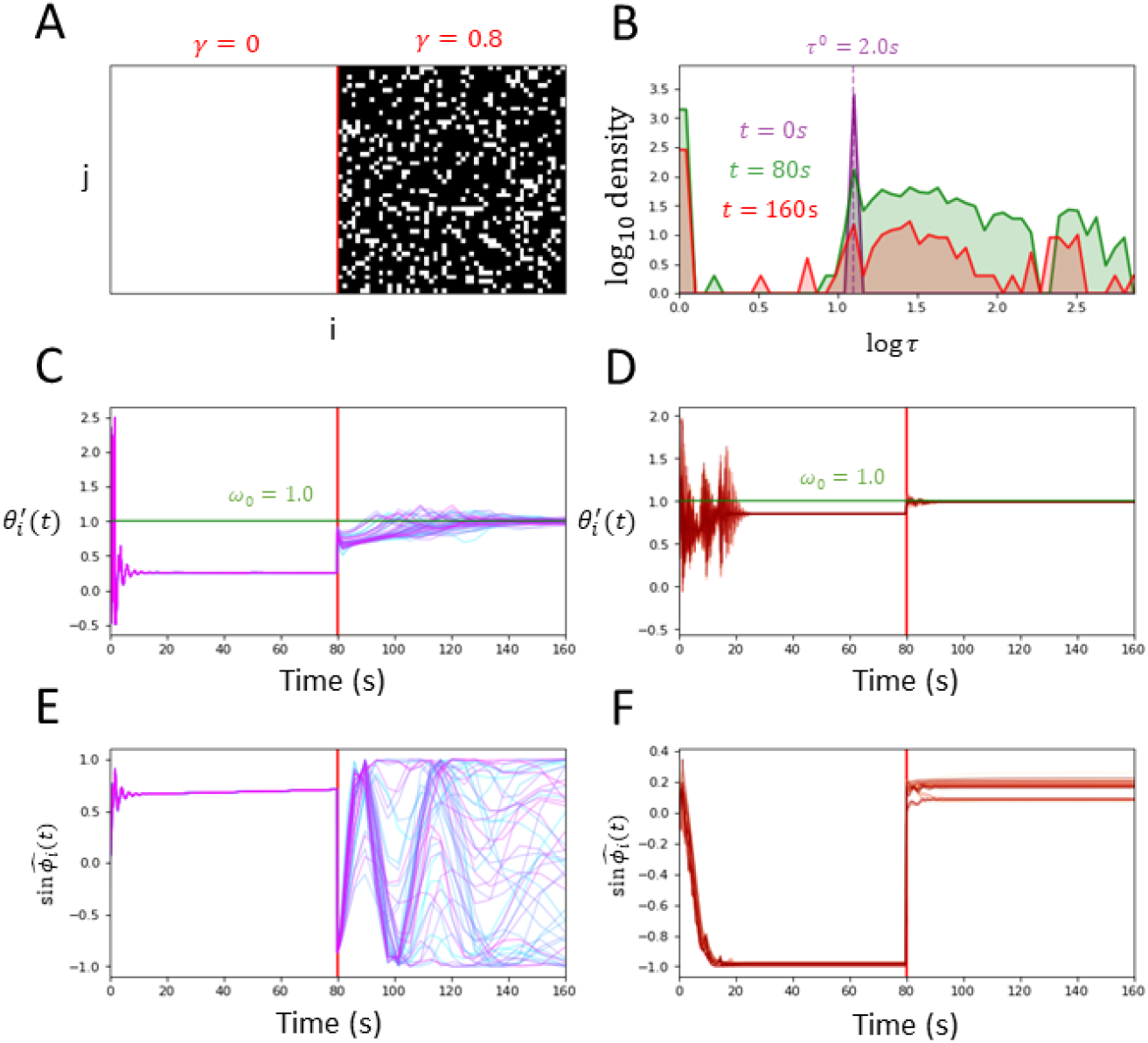
Comparing resilience against injury between plastic and non-plastic delays. Injury towards the connection topology *a_ij_* is introduced at *t_inj_* = 160s (red line). A. Plots of the connection matrix *A* = (*a_ij_*) before injury (*γ* = 0) and after injury (*γ* = 0.8), with *a_ij_* = 1,0 indicated in white, black respectively. B. The log histogram plots of delays at initial time *t* = 0*s* (purple), midtime before injury *t* = 160s (green), and at the end time following injury *t* = 320*s* (red). The delays become distributed away from *τ_ij_* = *τ*^0^ to either some largely varying equilibrium delays 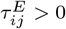 or decay to 0. Following injury, there are fewer delays available to stabilize the synchronous network. C, D. Plots of derivatives 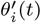 over time, without and with plasticity respectively. Both networks entrain in frequency pre-injury. Following injury, both networks entrain to a new frequency closer to *ω*_0_. E, F. Plots of 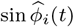 over time, without and with plasticity respectively. Following injury, the oscillators with plastic delays are able to coherently phase lock within close proximity to each other, while the network without plastic delays remain in an non-convergent state. The parameters used were *N* = 50, *ε* = 0.01, *α_τ_* = 1.0, *g* = 1.5, *κ* = 80, *ω*_0_ = 1.0, and *τ*^0^ = 2.0*s*.

Following injury, from Fig. 6(E) the network without adaptive delays collectively falls out of phase. In contrast, Fig. 6(F) shows the network with adaptive delays demonstrating resilience against the injury as most oscillators are able to collectively phase-lock within close proximity to each other.

Fig. 7 examines the effect of gradually increasing the severity of injury towards the system’s global frequency 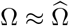 and its phase offset variance 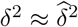. For each trial at injury *γ*, the same initial condition and parameters were used as in Fig. 6. Fig. 7(A) shows that connection loss generally leads to the system’s synchronization frequency Ω becoming closer to the natural frequency *ω*_0_. Fig. 7(B) plots the estimated phase standard deviation 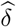 with respect to increasing injury. The network without plastic delays exhibits a significant loss in coherent synchrony with increasing 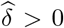 as more connections are lost. In contrast, the network equipped with adaptive delays persistently displays phase coherence with 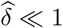 until higher injury levels 7.

**Figure 7.**
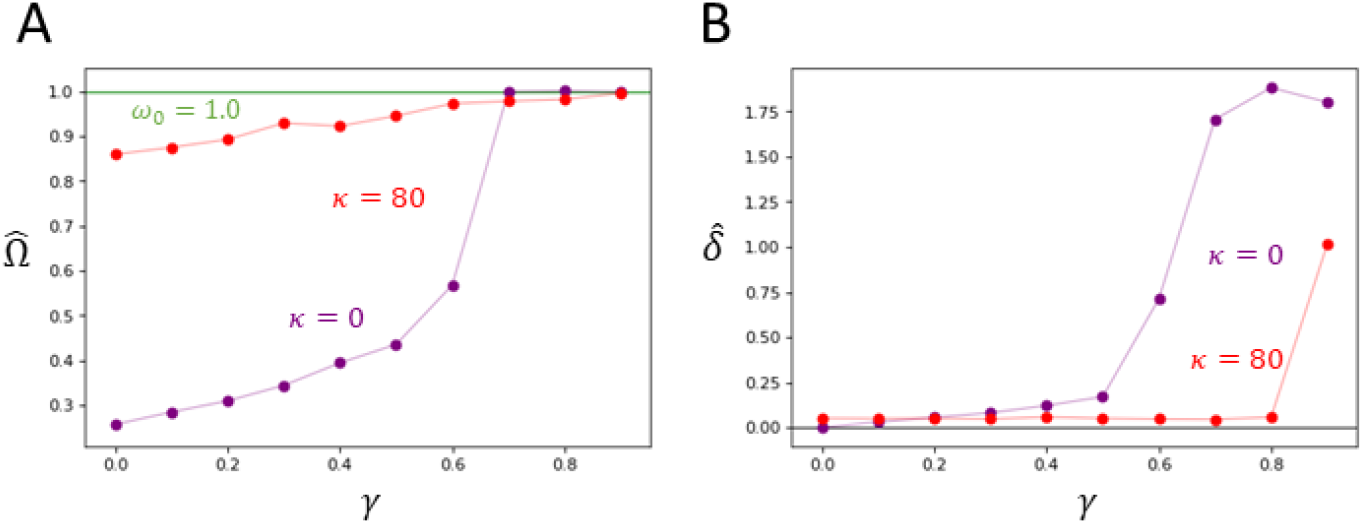
Comparing of post-injury network asymptotic behaviour with increasing injury between plastic and non-plastic delays. Each numerical trial was run over 320*s*, and all asymptotic values were evaluated by averaging over the final 16*s*. Injury was introduced at *t* = 160*s*. Both plots show trials with adaptive delays (red) and fixed delays (purple). A. Plot of post-injury asymptotic frequency 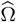 for trials with injury *γ* > 0. As *γ* increases, 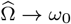. B. Plot of post-injury asymptotic standard deviation 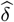 for trials with injury *γ* > 0. As *γ* increases, 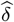 has significant increase for trials without plasticity, while 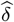 remains relatively small for trials with plasticity until *γ* = 0.9. The parameters used were *N* = 50, *ε* = 0.01, *α_τ_* = 1.0, *g* = 1.5, *κ* = 80, *ω*_0_ = 1.0, and *τ*^0^ = 2.0*s*.

## 8 Discussion

Our goal was to provide a mathematical framework that captures the synchronizing properties of networks with adaptive delays. We sought to implement the activity-dependent property of myelinated connection delays by modifying the Kuramoto model as proposed in [9]. The focus was to determine whether such adaptive delays significantly improve the oscillatory system’s ability to become in-phase and to entrain to a global frequency. Given that this is the case, the results of the model’s study reinforces the proposition that myelin plasticity is essential in maintaining the synchrony in the developing or injured brain.

White matter plays a critical role in maintaining brain function through the coordination of neural dynamics across multiple temporal and spatial scales. Recent evidence has shown that through the action of glia, white matter properties evolve continuously in time. Specifically, conduction velocity within and across brain areas is adjusted to promote efficient neural signaling. While the mechanism remains poorly understood, the consequences of such plastic processes on brain dynamics and synchronization can be readily examined and characterized using simplified mathematical models.

To accomplish this, we here examined the influence of adaptive conduction delays on the synchronization of neural oscillators. We developed a repertoire of mathematical tools to better examine the stability of phase locked solutions. In theory, we derived implicit equations for the global frequency Ω and eigenvalues 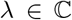 that provide a rigorous criterion for the stability around the synchronous state in two-dimensional and large-dimensional settings. Based on our model, flexibility in the white matter structure introduces an additional corrective dynamic next to the phase interactions that can further drive the network’s phase alignment. Higher stability with adaptive delays was demonstrated as the Kuramoto model had higher resilience against injury perturbations. However, adaptive delays improve the system’s synchronous features only when the delays adjust with a sufficiently high degree of plasticity, as represented by the plasticity gain *κ*.

There are many limitations in the prototypical model we used and its corresponding results. Myelination is bound by many physiological constraints, some of which remain uncertain [4]. It is established that white matter restructures itself in response to ongoing neural activity [8]. We primarily incorporated this fact in our plasticity rule in a manner that promotes local synchrony. Indeed, each connection delay changes at a rate proportional to the sine of the oscillator’s phase difference. This rule remains a tentative construction, as more research is needed to develop more biological relevant models in activity-dependent myelination. In addition, the use of phase oscillators to model local neural dynamics remain limited and is relevant mostly in the context of large-scale neural systems. In our analysis, we relied heavily on i.i.d. parametrical frameworks in order to establish our *N*-limit approach, which may not be feasible as network elements are correlative in nature.

Moving forward, we hope to build upon our analysis alongside newly found experimental results pertaining to myelin. Despite its shortcomings, the mathematical approaches used and its results can potentially be applied to more complex, biological relevant models. The conduction delays 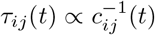 can be alternatively modelled with respect to a system of adaptive conduction velocities *c_ij_*(*t*). In the realm of temporal equations, other parametric avenues have yet to be explored. For instance, the delays *τ_ij_*(*t*) can exhibit slow convergence by setting the rate constant *α_τ_* ≪ 1. The aforementioned concepts are some proposed examples that may further lead to uncharted dynamics in the scope of neurocomputational models.

## Appendix A: Existence, uniqueness, and linearization

We sketch out the arguments needed to justify the existence, uniqueness, and stability analysis of our proposed Kuramoto model with adaptive delays. Our model can be expressed as an *N* + *N*^2^ system of oscillators 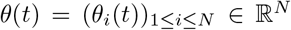 and delays 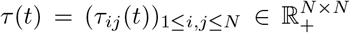, where 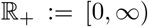, defined by the state-dependent Functional Differential Equations

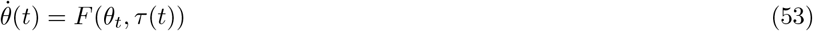

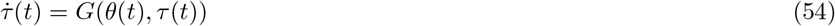

where 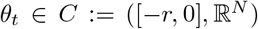 defined by *θ_t_*(*s*) = *θ*(*t* + *s*), *s* ∈ [−*r*, 0], and *C* is the space of all continuous functions from interval [−*r*, 0] for some *r* > 0. Here, 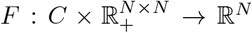 is the function whose ith co-ordinate *F_i_*(*θ_t_, τ*(*t*)) is given by Eq. (1) and 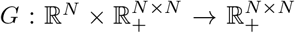 is the function whose *ij*th co-ordinate *G_ij_*(*θ_i_*(*t*), *θ_j_*(*t*), *τ_ij_*(*t*)) is given by Eq. (3). We note that from Eq. (3), each delay solution *τ*(*t*) is bounded for all *t* ≥ 0. This is clear since given *τ_ij_*(*t*) > *τ*^0^ + *κ*, it must hold that 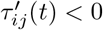. Hence, it suffices to define the space *C* with any *r* > *τ*^0^ + *κ*.

Denote *C*^1^ ⊂ *C* to be the subspace of all continuously differentiable functions. Let *θ*_0_(*s*) = *φ*(*s*) for some *C*^1^ initial function 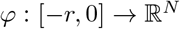 and 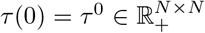 be the initial delays. Since Eq. (54) is a system of Ordinary Differential Equations, *τ*(*t*) is independent of initial values on *t* < 0. Hence, we represent an initial function for *τ*(*t*) with an initial point 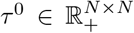. Existence and uniqueness of solution (*θ*(*t*), *τ*(*t*)) follows from theorems in [18] given that both *F, G* are continuously differentiable functions over 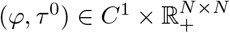. The *i*th co-ordinate *F_i_*(*φ, τ*^0^) is a linear combination of terms 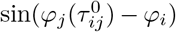, which is continuously differentiable over *φ* ∈ *C*^1^. *G* is continuously differentiable given the Heaviside function *H*(*τ*) is smooth. The Heaviside function used throughout the paper is constructed as follows. Setting some arbitrarily small *ε* > 0, we define the smooth mollifier function 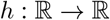 by

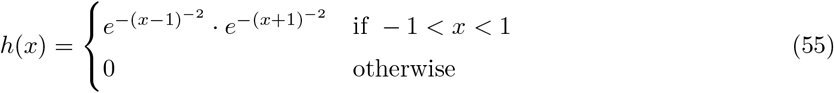

Then, our smooth Heaviside function *H*(*τ*) is defined as

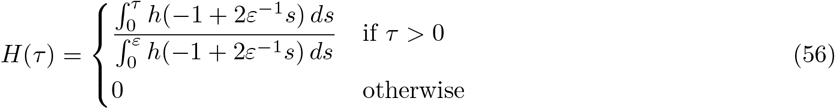

Then, each co-ordinate *G_ij_* is smooth with respect to terms *φ_i_, φ_j_, τ*^0^.

Linearization and stability of our system around the synchronized state (Ω, *ϕ*) can be justified as follows. We can express Eqs. (53, 54) under perturbation terms (*ε*(*t*), *η*(*t*)) of (*θ*(*t*), *τ*(*t*)) defined by Eqs. (7, 8). That is, consider the autonomous equations

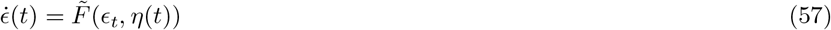

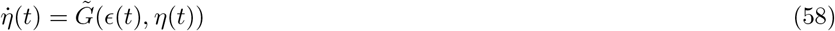

where the *i*th co-ordinate of 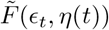 is given by Eq. (10) and 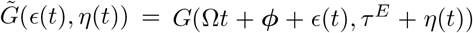. Then, if (Ω, *ϕ*) satisfies Eq. (5), 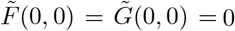. Hence, the stability of (*θ, τ*) around (Ω, *ϕ*) depends on the linearization of 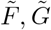 around (0,0). One can compute that the corresponding linearized system 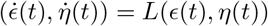 is given by Eqs. (9, 11). To prove that this linearization is valid and provides the eigenvalue stability criteria det *M*_λ_ = 0, we can replicate the proof for Theorem 2.1 in [20]. Indeed, 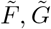 are composed of Lipschitz continuous terms that allow the steps to be valid for our perturbation Eqs. (57, 58).

## Appendix B: Derivation of N-limit equations

Here, we justify the limit steps taken in Sect. 5 to derive the limit-*N* equations for global frequency Ω and stability eigenvalues λ around the synchronous state. First, denote 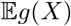 as the expectation of random variable *g*(*X*). That is, given *X* has distribution *ρ*(*x*),

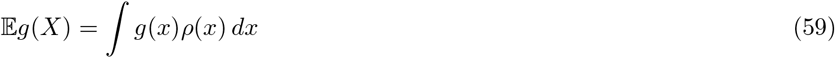

The main proposition we applied is as follows.

### Theorem 1

*Let* (*v_k_*)_*k*≥1_ *be a bounded sequence of real numbers, and X be a random variable such that* 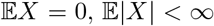. *Suppose* (*X_k_*)_*k*≥1_ *is an i.i.d. sequence of random variables such that each X_i_ has the same distribution as X. Then*,

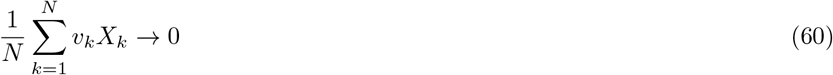

*almost surely as N* → ∞.

*Proof* For some *M* > 0, |*v_k_*| < *M* for all *k*. By the classic Strong Law of Large Numbers,

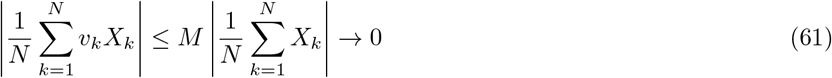

almost surely as *N* → ∞.

What directly follows from Theorem 1 is the following limit result, which we utilize to derive the limit-N equations for the global frequency Ω and the stability eigenvalues λ.

### Corollary 1

*Let* (*v_k_*)_*k*≥1_ *be a bounded sequence in* 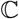, *and X be a random variable such that* 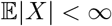. *Suppose* (*X_k_*)_*k*≥1_ *is an i.i.d. sequence of random variables such that each X_i_ has the same distribution as X. Then*,

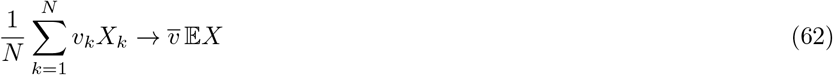

*where* 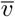 *is the well-defined asymptotic sample mean of the coefficients:*

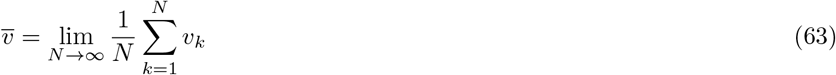

*In particular, if v*: 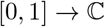 *is a continuous function and v_k_* = *v*(*k/N*), *then*

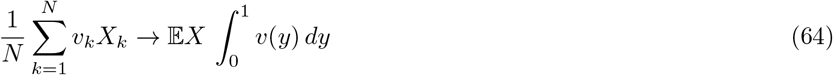

*Proof* This immediately follows by applying Theorem 1 to the i.i.d. sequence 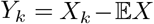. It remains to show that 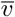 exists. Denoting 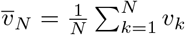 as the sample mean among the first *N* coefficients with |*v_k_*| ≤ *M* for all *k*, we show that 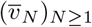 is a Cauchy sequence. We have:

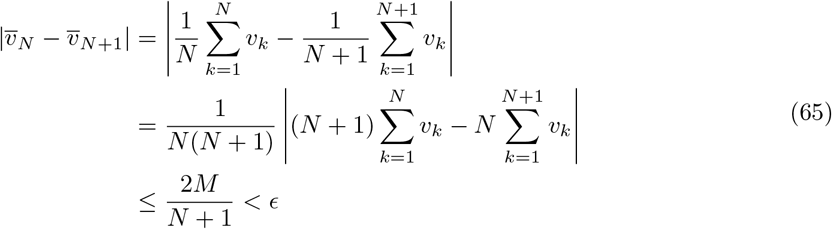

for sufficiently large *N*. Hence, the limit 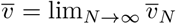 exists. In the case where *v_i_* = *v*(*i/N*), it is clear that 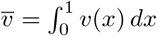 by taking Riemann sums.

We applied Corollary 1 to random sequences of the form *X_k_* = *f* (Δ_*k*_, *τ*^0^) where Δ_*k*_ is Gaussian distributed, and *τ*^0^ is the constant baseline lag. The eigenfunction coefficients *v_k_* are from ansatz *ε_k_*(*t*) = *v_k_e*^λ*t*^ as *N* → ∞.

## Appendix C: Numerical setup

All numerical simulations in Sects. 6, 7 were done using MATLAB’s ddesd function. We have a system of delay differential equations for the phases *θ_i_*(*t*), *i* ≤ *N* given by Eq. (1) and an *N*^2^ system of ordinary differential equations for the state-dependent delays *τ_ij_*(*t*), 1 ≤ *i, j* ≤ *N* given by Eq. (3). The initial data before starting time *t* = 0 are as follows. We have a history function 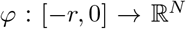 for the phases, such that *θ_i_*(*t*) = *φ_i_*(*t*) for all *t* ≤ 0. We considered only the simple case where delays *τ_ij_*(*t*) have a unique baseline lag 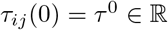. As shown in Appendix A, it is sufficient to consider *C*^1^ initial phase functions *φ*(*t*) to ensure unique solutions. However, we are not guaranteed to have reasonably behaving solutions unless we have continuity of *θ*′(*t*) at *t* = 0 [19]. This is also a necessary condition for providing accurate numerical simulations, as interpolation steps in `ddesd` rely on *θ*′(*t*) being continuous everywhere.

For our numerical experiments in Sects. 6, 7, we consider the linear initial functions 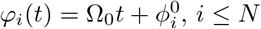 with respect to initial frequency 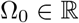 and phase offsets 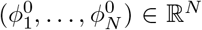. As discussed above, we require that our linear initial function must satisfy the necessary condition Eq. (49). Given any initial *C*^1^ function *φ*(*t*) for *θ*(*t*), we define the modified *C*^1^ function 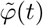 of *φ*(*t*) that satisfies the necessary condition as follows. We can obtain the modified slope 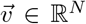 by imposing the condition on *φ*(*t*). That is,

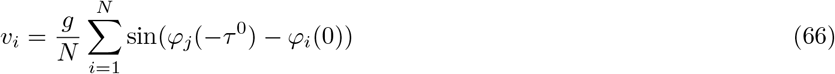

Then, for each *i* we define the cubic polynomial 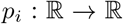 that interpolates *φ*(*t*) between *t* = 0, −*τ*^0^ using the modified slope. That is, *p_i_*(*t*) = *φ_i_*(*t*) at *t* = 0, −*τ*^0^ and 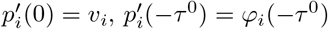. Then, the modified initial function 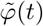 is given by

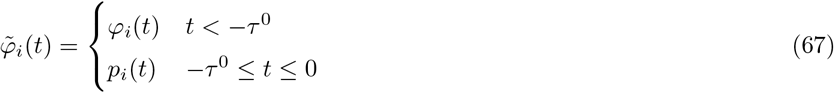

for *i* ≤ *N*. All numerical trials were conducted using the corresponding modified functions 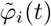 of 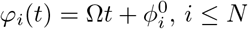.

## Abbreviations

i.i.d.: independent and identically distributed

## Ethics approval and consent to participate

Not applicable.

## Consent for publication

Not applicable.

## Availability of data and materials

The datasets generated and/or analysed during the current study are available in the github repository [27].

## Authors’ contributions

SHP and JL performed the research, did the analysis and wrote the manuscript.

## Acknowledgments

We would like to thank Adam Stinchcombe for his valuable feedback regarding the numerical methods.

## Footnotes

[1] We can represent any connection topology of the network as an integral operator *a*: [0, 1]^2^ → {0, 1} such that *a_ij_* = *a*(*i/N, j/N*), for which we find the eigenfunction 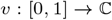 satisfying 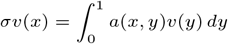 with respect to eigenvalue *σ*.

[2] Trials were done using other types of initial functions *φ_i_*(*i*), and yielded similar convergence results. However, the synchronization occurred at a slower rate.

## References

1. Strogatz SH. From Kuramoto to Crawford: exploring the onset of synchronization in populations of coupled oscillators. Physica D: Nonlinear Phenomena. 2000;143(1):1 – 20.

2. Buzsáki G. Rhythms of The Brain; 2009.

3. Cabral J, Luckhoo H, Woolrich M, Joensson M, Mohseni H, Baker A, et al. Exploring mechanisms of spontaneous MEG functional connectivity: How delayed network interactions lead to structured amplitude envelopes of band-pass filtered oscillations. Neuroimage. 2013 12;90.

4. de Hoz L, Simons M. The emerging functions of oligodendrocytes in regulating neuronal network behaviour. BioEssays. 2015 01;37.

5. Monje M. Myelin Plasticity and Nervous System Function. Annual Review of Neuroscience. 2018 07;41:61–76.

6. Bechler M, Swire M, Ffrench-Constant C. Intrinsic and adaptive myelination - a sequential mechanism for smart wiring in the brain. Developmental neurobiology. 2017 8;78(2):68–79.

7. G Almeida R, A Lyons D. On Myelinated Axon Plasticity and Neuronal Circuit Formation and Function. The Journal of Neuroscience. 2017 10;37(42):10023–10034.

8. Douglas Fields R. A new mechanism of nervous system plasticity: Activity-dependent myelination. Nature Reviews Neuroscience. 2015 11;16:756–767.

9. Pajevic S, Basser PJ, Fields RD. Role of myelin plasticity in oscillations and synchrony of neuronal activity. Neuroscience. 2014;276:135 – 147. Secrets of the CNS White Matter.

10. Breakspear M, Heitmann S, Daffertshofer A. Generative Models of Cortical Oscillations: Neurobiological Implications of the Kuramoto Model. 2010;4:190.

11. Izhikevich EM. Phase models with explicit time delays. Phys Rev E. 1998 Jul;58:905–908.

12. Yeung MKS, Strogatz SH. Time Delay in the Kuramoto Model of Coupled Oscillators. Phys Rev Lett. 1999 Jan;82:648–651.

13. Petkoski S, Jirsa V. Transmission time delays organize the brain network synchronization. Philosophical Transactions of the Royal Society A: Mathematical, Physical and Engineering Sciences. 2019 09;377:20180132.

14. Petkoski S, Palva JM, Jirsa V. Phase-lags in large scale brain synchronization: Methodological considerations and in-silico analysis. PLOS Computational Biology. 2018 07;14:e1006160.

15. Ponce-Alvarez A, Deco G, Hagmann P, Romani GL, Mantini D, Corbetta M. Resting-State Temporal Synchronization Networks Emerge from Connectivity Topology and Heterogeneity. PLoS computational biology. 2015 02;11:e1004100.

16. Dhamala M, Jirsa VK, Ding M. Enhancement of Neural Synchrony by Time Delay. Phys Rev Lett. 2004 Feb;92:074104.

17. Fries P. A mechanism for cognitive dynamics: neuronal communication through neuronal coherence. Trends in Cognitive Sciences. 2005;9(10):474 – 480.

18. Hale JK, Lunel SMV. Introduction to Functional Differential Equations. Springer-Verlag; 1993.

19. Hartung F, Krisztin T, Walther HO, Wu J. Chapter 5 Functional Differential Equations with State-Dependent Delays: Theory and Applications. 2006;3:435 – 545. Available from: http://www.sciencedirect.com/science/article/pii/S187457250680009X.

20. Cooke KL, Huang W. On the Problem of Linearization for State-Dependent Delay Differential Equations. Proceedings of the American Mathematical Society. 1996;124(5):1417–1426.

21. Papachristodoulou A, Jadbabaie A. Synchronization in Oscillator Networks: Switching Topologies and Non-homogeneous Delays. Proceedings of the 44th IEEE Conference on Decision and Control. 2005;p. 5692–5697.

22. Earl MG, Strogatz SH. Synchronization in oscillator networks with delayed coupling: A stability criterion. Phys Rev E. 2003 Mar;67:036204.

23. Atay F. Distributed delays facilitate amplitude death of coupled oscillators. Physical Review Letters, v91 (2003). 2003 01;91.

24. Cooke D. On Zeroes of Some Transcendental Equations. Funkcialaj Ekvacioj. 1986;p. 77–90.

25. Russell RD, Shampine LF. Existence of Eigenvalues of Integral Equations. SIAM Review. 1971;13(2):209–219. Available from: https://doi.org/10.1137/1013038.

26. Bells S, Lefebvre J, Prescott SA, Dockstader C, Bouffet E, Skocic J, et al. Changes in white matter microstructure impact cognition by disrupting the ability of neural assemblies to synchronize. Journal of Neuroscience. 2017;Available from: https://www.jneurosci.org/content/early/2017/07/25/JNEUROSCI.0560-17.2017.

27. Park SH. Kuramoto Plastic Delay;. Available from: https://github.com/parkseo7/KuramotoPlasticDelay.git. Accessed March 27 2020.

